# High-throughput spheroid profiling reveals BMP-driven rewiring of glioma cell death responses

**DOI:** 10.64898/2026.09.10.750657

**Authors:** Weaverly Colleen Lee, Jennifer J. Salinas, Aastha Gautam, Dimitri Cadet, Matei A. Banu, David A. Nathanson, Scott J. Dixon

## Abstract

The mechanisms regulating glioma cell death are poorly understood. Here, we developed a high-throughput method to study cell death in patient-derived glioblastoma (GBM) and diffuse intrinsic pontine glioma (DIPG) spheroids. Using this method, we systematically profiled how extracellular ligands modulate compound-induced cell death. We find that bone morphogenetic protein 2 (BMP2) and BMP4 potently rewire cell death sensitivity. These ligands suppress killing by standard-of-care DNA alkylating agents and kinase inhibitors by inhibiting cell cycle progression. Simultaneously, BMP2/4 prime spheroids for lipid-dependent necrosis (LiDN), a palmitate-dependent form of non-apoptotic cell death that can be triggered by the clinical drug candidate tegavivint. Activating mutations in the BMP receptor ACVR1, found in ∼25% of DIPG tumors, are sufficient to prime cells for LiDN in the absence of BMP ligand. Together, these findings identify a cell death switch that can be activated in glioma cells by BMP signaling.

## Main text

Glioblastoma (GBM) is a highly treatment-refractory brain cancer where cytotoxic therapies fail to produce durable tumor responses^1–3^. Standard-of-care agents such as the DNA alkylating drug temozolomide can engage apoptosis^2–5^. Emerging targeted therapies, including inhibitors of the epidermal growth factor receptor (EGFR) and phosphatidylinositol 3-kinase (PI3K), may also depend on apoptotic pathways for full killing efficacy^6–11^. However, treatment-naïve GBM can exist in a slow-cycling, therapy-resistant state^12^, or acquire genetic alterations that inhibit apoptosis^5, 13, 14^. Strategies to engage non-apoptotic cell death in brain cancer are potentially attractive^15^, but it is unclear how best to do so.

Most high-throughput cell death analyses use established cell lines growing in simple adherent or suspension conditions. However, clinically relevant brain cancer phenotypes are best preserved in three-dimensional patient-derived spheroids^16^. Analyzing cell death in spheroid models is not straightforward; spheroid drug responses are typically studied using assays that measure cell viability instead of cell death^5, 17^. Results obtained using these methods can be confounded by altered metabolism, proliferative arrest and incomplete killing^18, 19^. To overcome these limitations, we developed an imaging-based method called High-throughput Analysis of Death in Spheroids (HADES) to directly quantify cell death in patient-derived spheroid models.

Cell death responses can be regulated by cell non-autonomous factors^20^. Brain tumors reside in microenvironments rich in extracellular ligands^17, 21–23^. Understanding how these ligands influence cell death responses could provide new insights into cell death regulation and potentially suggest new treatment strategies. Using HADES, we profiled GBM-relevant protein ligands for effects on compound-induced death. Bone morphogenetic protein 2 (BMP2) and BMP4 emerged as potent rewiring signals that acted through the canonical BMP receptor-SMAD pathway to suppress killing by DNA alkylating agents and kinase inhibitors while sensitizing cells to lipid-dependent necrosis (LiDN), a non-apoptotic cell death mechanism^24, 25^. We extended these findings to diffuse intrinsic pontine glioma (DIPG), where activating mutations are found in the BMP receptor *ACVR1* in ∼25% of tumors^26–29^. Together, these findings establish a scalable platform to directly profile cell death in patient-derived spheroid models and identify BMP signaling as a driver of a cell death sensitivity switch in brain cancer.

## RESULTS

### High-throughput analysis of spheroid cell death

We sought to examine compound-induced cell death in glioma, starting with GBM. Primary GBM cells grown as non-adherent gliomaspheres maintain tumor phenotypes but are difficult to use in high-throughput cell death assays^5, 16–18, 30^. To address this, we developed HADES, an imaging-based method to quantify cell death in gliomaspheres and other spheroid models. In this method, low-passage cells are engineered to express nuclear-localized mKate2 (denoted by the superscript “N”, e.g., GS025^N^) as a live-cell marker and cultured with the dead-cell stain SYTOX Green^19^. The loss of red fluorescence and gain of green fluorescence within each spheroid over time is used to infer cell death, which is reported as the spheroid death score (**Fig. 1a**).

**Figure 1.**
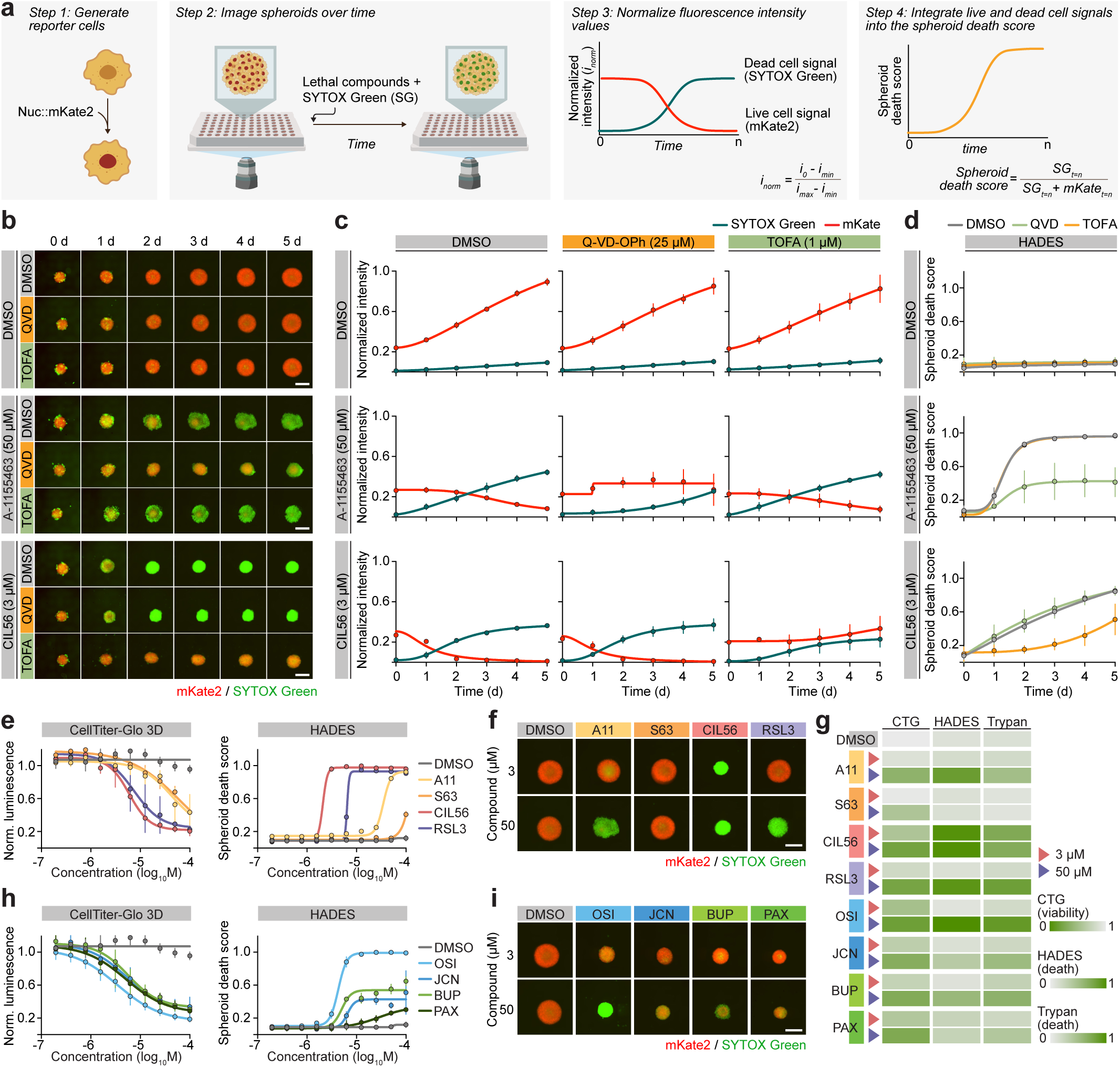
High-throughput analysis of gliomasphere cell death. a, Overview of the High-throughput Analysis of Death in Spheroids (HADES) method. b, Imaging of GS025^N^ gliomaspheres. c, Normalized fluorescence intensity values for mKate2 (live cell) and SYTOX Green (dead cell) reporters in treated GS025^N^ gliomaspheres. d, Spheroid death scores, calculated from data in (c). Higher values indicate more death within the gliomasphere. e, Comparison of HADES with CellTiter-Glo 3D (CTG) in spheroids treated with compounds targeting specific cell death pathways. A11: A-1155463, S63:S63845. f, GS025^N^ gliomasphere images. g, Heatmaps representing viability or cell death, measured by CTG, HADES or trypan blue staining. h, Comparison of HADES with CTG in spheroids treated with compounds targeting growth factor signaling pathways. OSI: osimertinib, JCN: JCN037, BUP: buparlisib, PAX: paxalisib. i, GS025^N^ gliomasphere images. Images in (b), (f) and (i) are representative of three experiments. Scale bars = 400 µm. Data in (c)-(e) and (h) represent mean ± s.d. from three independent experiments. Data in (j) summarize results from three independent experiments.

We used HADES to study cell death in patient-derived gliomaspheres from different molecular subtypes and transcriptional cell states^31^ (**Extended Data Fig. 1a-c, Extended Data Table 1,2**). To initially assess this method, GS025^N^ gliomaspheres were treated with the BCL2 like 1 (BCL2L1, BCL-xL) inhibitor A-1155463 (ref.^32^) or the LiDN inducer CIL56 (ref.^24, 25^). Both compounds caused loss of mKate2 signal and gain of SYTOX Green signal over time, and these effects were selectively blocked by the pan-caspase inhibitor Q-VD-OPh^33^ and the acetyl-CoA carboxylase inhibitor 5-(tetradecyloxy)-2-furoic acid (TOFA)^24, 25^, which inhibit apoptosis and LiDN, respectively (**Fig. 1b-d**). Thus, HADES detected both apoptotic and non-apoptotic cell death in gliomaspheres.

We compared HADES with CellTiter-Glo 3D (CTG), an ATP-based viability assay commonly used with spheroid models^5, 17^. GS025^N^ gliomaspheres were treated with the BCL-xL inhibitor A-1155463, the MCL1 inhibitor S63845, the LiDN inducer CIL56, or the ferroptosis inducer RSL3, each in a 10-point, two-fold dilution series. Dose-dependent differences in the lethal potency of these four agents were readily apparent using HADES but not CTG (**Fig. 1e,f**). For example, the higher potency of CIL56 versus RSL3 and A-1155463 versus S63845 were uniquely detected using HADES (**Fig. 1e,f**). These differences in cell death induction were confirmed using trypan blue staining, a distinct low-throughput cell death detection method (**Fig. 1g**). We next expanding our analysis to candidate GBM therapies: EGFR inhibitors JCN037 and osimertinib, and the PI3K inhibitors paxalisib and buparlisib^7–11^. All four kinase inhibitors uniformly reduced cell viability as determined using CTG. By contrast, clear differences in cell death were apparent between these kinase inhibitors when assessed using HADES or trypan blue staining, with osimertinib and buparlisib yielding greater cell death than JCN037 or paxalisib (**Fig. 1g-i**). All four kinase inhibitors limited gliomasphere growth over time and inhibited proliferation in adherent T98G^N^ cells (**Extended Data Fig. 1e,f**). These results suggested that HADES could effectively detect cell death and distinguish this from growth arrest in primary gliomaspheres.

### Extracellular ligands modulate GBM cell death

The brain environment contains dozens of extracellular ligands that can modulate cell fate^34^. We aimed to use HADES to systematically study the impact of individual extracellular protein ligands on drug-induced cell death (**Fig. 2a**). For this analysis we focused on 13 drugs or tools compounds that are or could potentially be used to treat GBM^9–11, 24, 25, 32, 35–42^, most of which we had already validated in our models (**Fig. 2b**). Potential modulatory ligands were identified as follows. RNA-sequencing coupled with published datasets^43–45^ were used to predict 1,919 potential ligand-receptor pairs in GS025 cells (**Extended Data Fig. 2a**). The clinical importance of each predicted ligand-receptor pair was then ranked by correlating ligand and receptor gene expression with GBM patient survival data from The Cancer Genome Atlas (**Fig. 2c,d** and **Extended Data Fig. 2a**). The 50 ligands with the highest correlations to patient survival that could be obtained commercially were selected for further analysis (**Extended Data Fig. 2b**).

**Figure 2.**
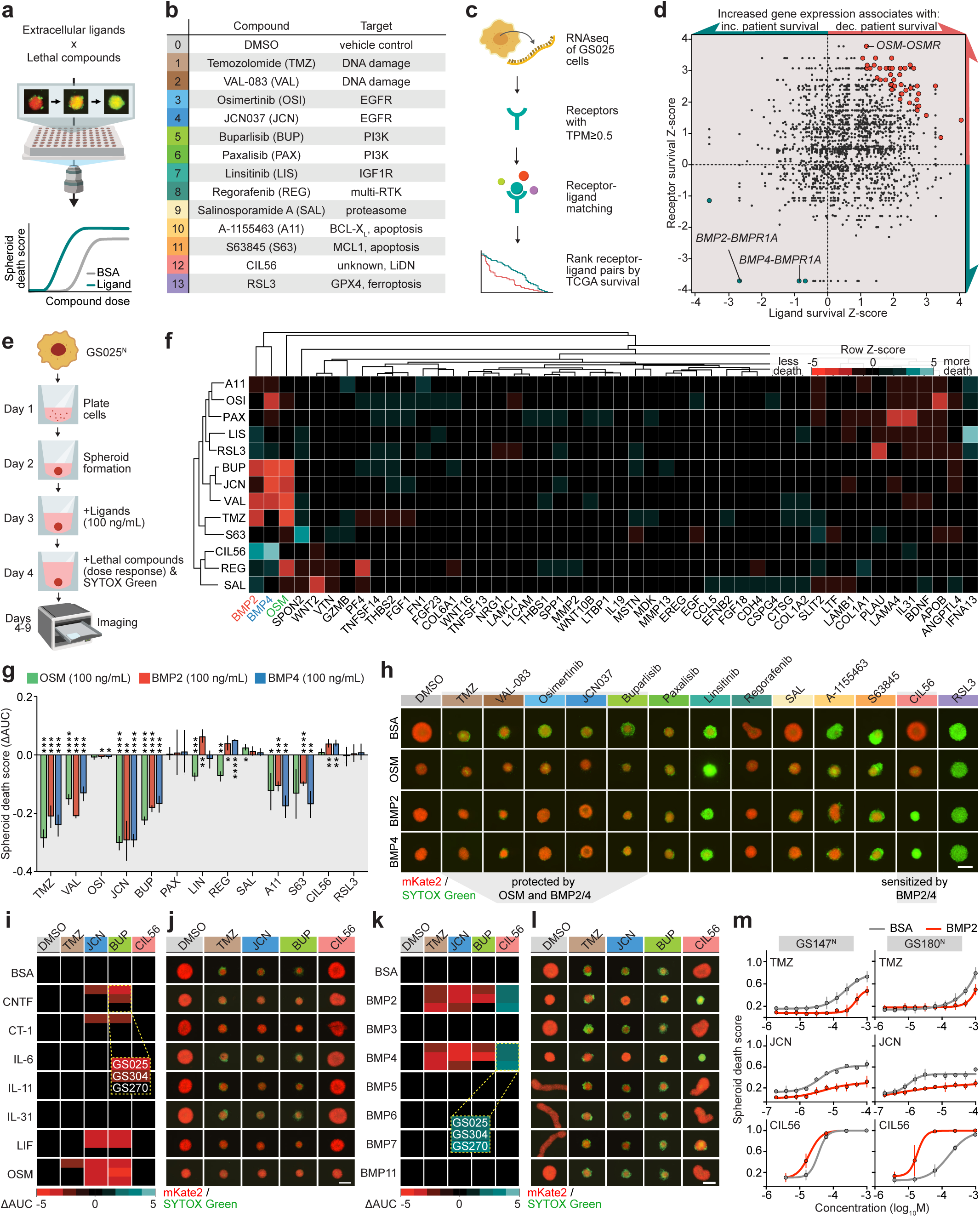
An extracellular ligand screen identifies modulators of cell death response. a, Schematic of ligand cell death modulator screen. b, Compounds used in the screen and primary targets where known. c, Overview of the bioinformatics workflow used to nominate ligands for screening. d, Combined survival Z-scores for predicted receptor-ligand pairs in GS025 gliomaspheres. Highlighted points denote the 50 pairs selected for screening. Red indicates association with decreased patient survival and blue indicates increased patient survival. Survival data were obtained from TCGA. e, Schematic of screening method. f, Heatmap summarizing the screen data. Colors reflect row-normalized Z-scores of the ΔAUC values (AUC_ligand_-AUC_BSA_). Red indicates ligand-mediated protection and blue indicates sensitization. g, Effects of OSM, BMP2 and BMP4 on GS025^N^ gliomasphere death. Ligands were used at 100 ng/mL. Asterisks indicate significance values from unpaired parametric t-tests comparing ligand-treated spheroids with matched BSA controls (*P ≤ 0.05, **P ≤ 0.01, ***P ≤ 0.001). h, Representative GS025^N^ gliomasphere images for data in panel g. i, Effects of OSM and related proteins on gliomasphere cell death. Ligands were added in a 10-point, 2-fold dilution series starting at 100 ng/mL and spheroid death was integrated across concentrations. Temozolomide was used at 1 mM, JCN037 at 10 µM, buparlisib at 8 µM and CIL56 at 800 nM. The experiment was performed in three patient-derived gliomasphere lines, each represented as a subrow for each condition. j, Representative images for data in panel i, showing GS025^N^ gliomaspheres treated with 100 ng/mL of ligands. k, Effects of BMP2, BMP4 and related proteins on gliomasphere death. Ligands were added in a 10-point, 2-fold dilution series starting at 100 ng/mL and spheroid death was integrated across concentrations. Temozolomide was used at 1 mM, JCN037 at 10 µM, buparlisib at 8 µM and CIL56 at 800 nM. The experiment was performed in three patient-derived gliomasphere lines, each represented as a subrow for each condition. l, Representative images for data in panel k, showing GS025^N^ gliomaspheres treated with 100 ng/mL of ligands. m, Effect of BMP2 on GS147^N^ and GS180^N^ gliomasphere cell death. BMP2 was used at 100 ng/mL. Data in (g) and (m) represent mean ± s.d. from three independent experiments. Heatmaps in (i) and (k) show mean values from three independent experiments. Images in (h), (j), and (l) are representative of three independent experiments. Scale bars = 400 µm.

Gliomaspheres are grown in a minimal, serum-free medium containing three factors necessary for basal proliferation: heparin (5 µg/mL), EGF (50 ng/mL), and fibroblast growth factor b (bFGF, 20 ng/mL)^15, 16^. The impact of individual ligands on cell death was assessed in GS025^N^ gliomaspheres growing in this basal medium, pretreated with each of the 50 modulatory ligands (100 ng/mL) for 24 h, and then challenged with the 13 lethal compounds in 10-point, two-fold dilution series. Cell death was measured using HADES for five days, generating 4,620 dose-response curves from 50,820 individual gliomasphere cell death measurements (**Fig. 2e**). Two ligands (LAMA1 and TGM2) appeared autofluorescent in the absence of compound treatment and were excluded from further analysis (**Extended Data Fig. 2c,d**).

HADES-derived cell death measurements were integrated across compound doses and over time, cell death modulation was determined in reference to untreated cells, and the results were initially summarized as a heatmap reflecting the ability of different ligands to enhance or suppress cell death (**Fig. 2f**). Overall, 39/50 ligands did not significantly modulate cell death in response to any lethal compound (Z-score < 2), and eight ligands modulated cell death in response to one lethal compound (**Fig. 2f**). Our attention was drawn to oncostatin M (OSM), bone morphogenetic protein 2 (BMP2) and BMP4, which consistently enhanced resistance to TMZ^46^, VAL-083, JCN037 buparlisib (**Fig. 2f**). We confirmed the potent resistance triggered by OSM, BMP2 and BMP4 towards TMZ, VAL-083, JCN037 and buparlisib in validation assays performed in GS025^N^ gliomaspheres, with these ligands having weaker or no effects on the dose-dependent lethality of other agents (**Fig. 2g-h, Extended Data Fig. 2e**). While OSM enhanced resistance to JCN037 and buparlisib in two other patient-derived gliomasphere lines (GS304^N^, GS270^N^), the ability of OSM to confer resistance to TMZ appeared restricted to GS025^N^ (**Fig. 2i**). Other ligands in the same family as OSM^47^, like leukemia inhibitory factor (LIF) and ciliary neurotrophic factor (CNTF), also increased resistance to JCN037 and buparlisib, but with weaker and more gliomasphere line-restricted effects (**Fig. 2j**).

Unlike OSM, BMP2 and BMP4 (BMP2/4) more consistently resulted in resistance to TMZ, JCN037 and buparlisib while simultaneously and uniquely enhancing sensitivity to the LiDN inducer CIL56 across patient-derived gliomasphere lines (**Fig. 2k** and **Extended Data Fig. 2e**). The effects of BMP2/4 on cell death were dose-dependent and not recapitulated by BMP3 or other BMP family ligands^48^ (**Fig. 2k,l, Extended Data Fig. 2f,g**). BMP2 also modulated compound-induced cell death in two additional gliomasphere lines, consistent with a generalizable effect that did not correlate with patient sex, common GBM mutations, or transcriptional cell state^31^ (**Fig. 2m, Extended Data Table 1**). Notably, BMP2 treatment did not modulate cell death in adherent T98G^N^ GBM cells despite biochemical evidence for BMP signaling pathway activation (**Extended Data Fig. 2h,i**). Thus, BMP2/4 appeared to rewire cell death sensitivity specifically in patient-derived gliomaspheres.

### BMP2/4 modulate cell death via BMPR-SMAD signaling

BMP2/4 signal through heterodimeric receptor complexes comprising the type I BMP receptors BMPR1A or BMPR1B and the type II receptor BMPR2^48^ (**Fig. 3a**). The original motivation for including BMP2 and BMP4 in our cell death modulation analysis had included the observation that *BMPR1* expression correlated with GBM patient survival (**Fig. 2d**), and we confirmed from single nuclear RNA sequencing data^49^ that human GBM tumor samples express *BMPR1A*, *BMPR1B*, and *BMPR2*, with increased BMP receptor expression during progression from primary to recurrent tumors (**Extended Data Fig. 3b**). To test whether BMP2/4 acted via BMPRs to modulate cell death, we combined genetic and pharmacological perturbations with HADES. *BMPR1A* silencing in GS025^N^ gliomaspheres using short hairpin RNA (shRNA) did not alter compound-induced cell death responses, likely due to incomplete protein loss or functional redundancy with BMPR1B (**Extended Data Fig. 4a-c** and **Extended Data Table 3**). We therefore used the small molecule inhibitor LDN193189 (ref.^50^) to block type I BMP receptor function and test pathway dependence. In GS025^N^ gliomaspheres, BMP2/4, but not BMP3, increased SMAD1/5/8 phosphorylation and this was abolished by LDN193189 co-treatment (**Fig. 3b**). LDN193189 also blunted the ability of BMP2 to modulate compound-induced cell death (**Fig. 3c,d** and **Extended Data Fig. 4d**). Similar results were obtained using LDN193189 in GS304^N^ and GS270^N^ gliomaspheres and in GS025^N^ gliomaspheres treated with two additional inhibitors of type I BMP receptors, DMH1 and dorsomorphin^51, 52^ (**Extended Data Fig. 4e-l**).

**Figure 3.**
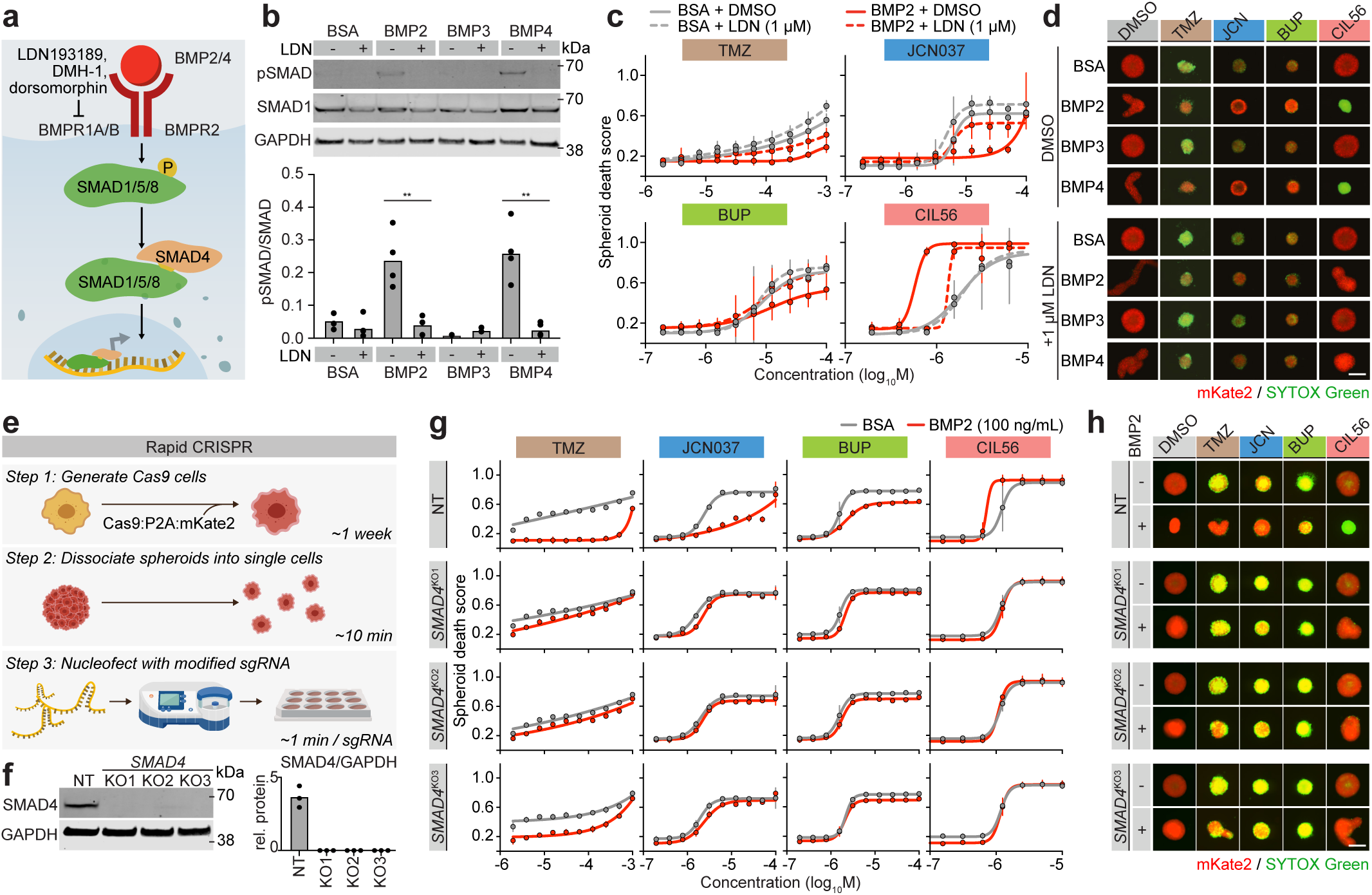
BMP2 and BMP4 modulate cell death through SMAD signaling. a, Schematic of the BMP2/4 signaling pathway and pharmacological inhibitors used in this study. b, Immunoblot analysis of SMAD1/5/8 phosphorylation (pSMAD) following LDN193189 (LDN; 1 µM) treatment. Blots are representative of four independent experiments. Asterisks indicate significance values from unpaired parametric t-tests comparing DMSO- and LDN-treated gliomaspheres (*P ≤ 0.05, **P ≤ 0.01, ***P ≤ 0.001). c, Spheroid death quantified by HADES analysis of GS025^N^ gliomaspheres treated with Temozolomide (TMZ), JCN037 (JCN), buparlisib (BUP) ± ligand ± LDN. d, Representative GS025^N^ gliomasphere images for data in panel c. TMZ was used at 500 µM, JCN at 12.5 µM, BUP at 12.5 µM, and CIL56 at 1.25 µM. e, Schematic of the rapid CRISPR (rCRISPR) method. f, Immunoblot validation of rCRISPR targeting of *SMAD4*. Blots are representative of three independent experiments; quantification is shown in the adjacent panel. g, Spheroid death quantified by HADES analysis of GS025^R^ gliomaspheres, with and without rCRISPR targeting of SMAD4. h, Representative GS025^R^ gliomasphere images for data in panel g using compound concentrations described in panel d. All ligand treatments = 100 ng/mL. Data in (c) and (g) represent mean ± s.d. from three independent experiments. Images in (d) and (h) are representative of three experiments. Scale bars = 400 µm.

BMPR1A/B phosphorylate SMAD1/5/8, transcription factors which complex with SMAD4 to regulate gene expression^48, 53^. In GS025^N^ cells, partial knockdown of *SMAD4* or overexpression of a dominant-negative *SMAD4^I500V^* mutant^54^ did not affect the lethality of TMZ, JCN037, or buparlisib, but attenuated the effects of BMP2 on CIL56-induced LiDN (**Extended Data Fig. 5a-f**). However, since shRNA and dominant-negative approaches may not completely inactivate SMAD4 function, we sought to generate *SMAD4* knockout cells. Conventional lentiviral-based CRISPR methods in primary GBM cells are slow and cumbersome^15^. Instead, GS025 cells stably expressing nuclear-localized Cas9 and mKate2 (denoted by the superscript “R”, i.e., GS025^R^) were directly nucleofected with chemically stabilized sgRNAs^55^ (**Fig. 3e**). This approach, hereafter referred to as rapid CRISPR (rCRISPR), yielded three *SMAD4* gene-disrupted (“knockout”, KO) polyclonal lines within one week (**Fig. 3f**). All three GS025^R^ *SMAD4^KO^* gliomasphere lines, but not a control line, were resistant to the effects of BMP2 on compound-induced cell death (**Fig. 3g,h**). Thus, canonical BMPR-SMAD signaling appeared to be required for BMP2 to rewire cell death responses in gliomaspheres.

### BMP2/4 arrest proliferation to protect from alkylating agents and kinase inhibitors

BMP2/4 consistently suppressed the lethality of TMZ, VAL-083, JCN037 and buparlisib in gliomaspheres (**Fig. 2g,k**). These compounds are reported to induce apoptosis in some settings^3–6^. We therefore initially hypothesized that BMP2/4 signaling might protect gliomaspheres by reducing apoptotic priming^56^. However, BH3 profiling showed the opposite: BMP2 increased gliomasphere apoptotic priming as measured by BIM-induced cytochrome c release (**Fig. 4a**). Consistent with heightened apoptotic sensitivity, BMP2-treated GS025^N^ gliomaspheres were sensitized to a combination of the BCL2-family inhibitors A-1155463 and S63845 (A11+S6), as determined using HADES (**Fig. 4b, Extended Data Fig. 6a**). The A11+S6 combination also increased caspase 3/7 processing and activity in these cells (**Fig. 4c,d**). By contrast, TMZ, VAL-083, JCN037 and buparlisib did not substantially increase caspase 3/7 activity in GS025^N^ gliomaspheres and cell death induced by these agents was not blocked by the pan-caspase inhibitor Q-VD-OPh (**Extended Data Fig. 6b**). Thus, BMP2/4 appeared to reduce sensitivity to the non-apoptotic effects of alkylating agents and kinase inhibitors.

**Figure 4.**
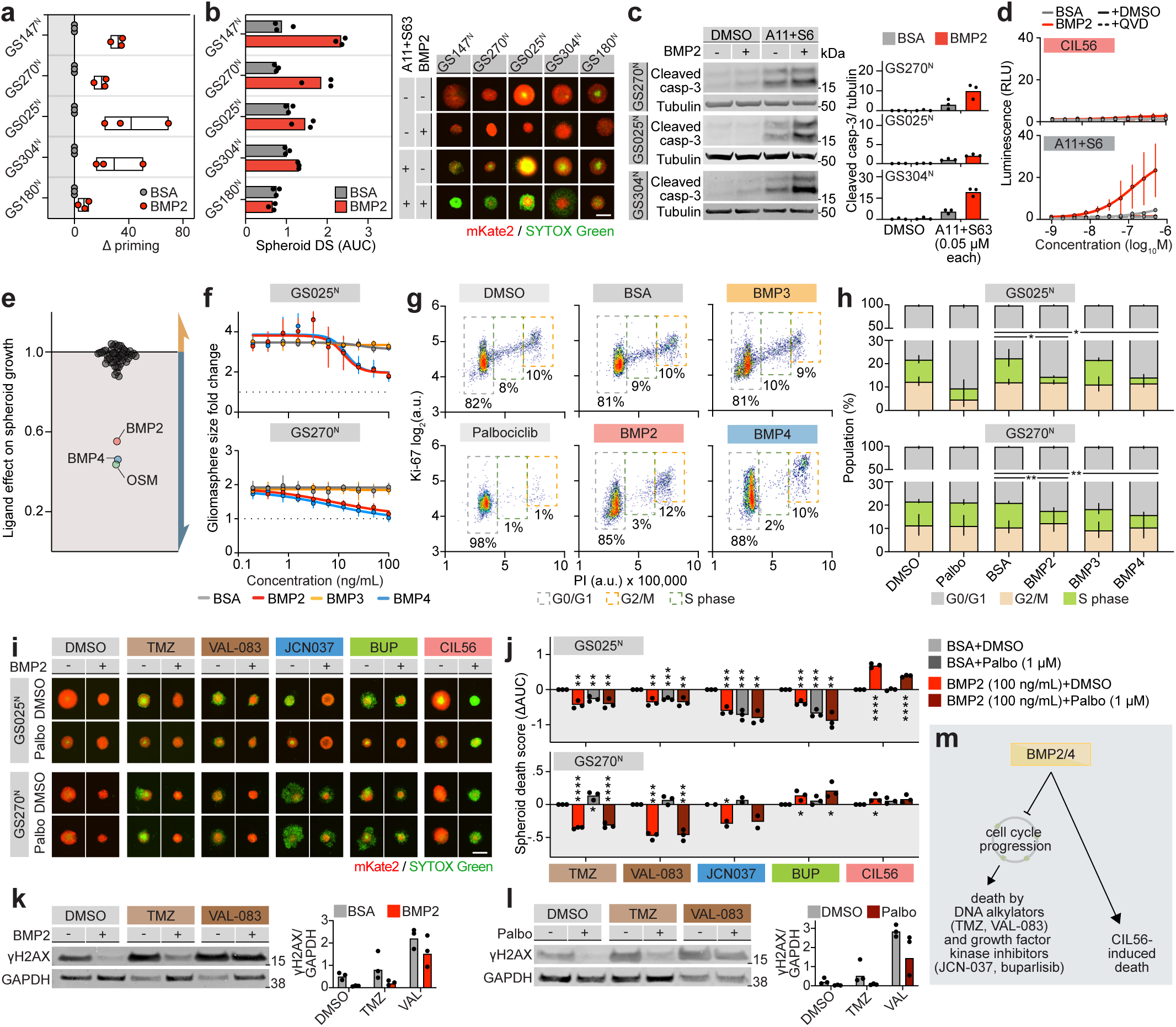
BMP2 and BMP4 decrease proliferation and death response to DNA alkylating agents and kinase inhibitors. a, Apoptotic priming determined using BH3 profiling with the BIM peptide. Δ priming = priming_BMP2_ - priming_BSA_. b, Cell death determined by imaging of gliomaspheres treated with A-1155463 (A11) and S63845 (S63). Representative images show gliomaspheres treated with 30 nM of each inhibitor; bar graphs summarize AUC values. c, Protein abundance determined by immunoblotting. BMP2 was used at 100 ng/mL, A-11 at 25 nM and S63 at 25 nM. d, Caspase-3/7 activity in GS025^N^ gliomaspheres determined by luminescence-based assay. A11 was used at 25 nM, S63 at 25 nM. e, Effects of the 50 screened ligands on GS025^N^ spheroid growth as quantified by imaging. f, Dose-dependent effects of BMP2 and BMP4 on GS025^N^ and GS270^N^ spheroid growth. Data are mean ± s.d. from three independent experiments. g, Representative flow cytometry cell cycle profiles of GS025^N^ gliomaspheres. Grey boxes indicate cells in G0/G1, green in S-phase, yellow in G2/M. h, Quantification of cell cycle distributions corresponding to data in panel c. Data are mean ± s.d. from three independent experiments. i, Spheroid death quantified by HADES analysis of gliomaspheres. Palbo, palbociclib. ΔAUC values (AUC_treated_ - AUC_BSA+DMSO_) summarize data from three independent experiments. j, Representative images of GS025^N^ and GS270^N^ gliomasphere for data in panel e. k, Immunoblot analysis of γH2AX in GS025^N^ cells following BMP2 treatment (100 ng/mL). l, Immunoblot analysis of γH2AX in GS025^N^ cells following Palbo treatment (1 µM). m, Model summarizing BMP2/4 effects on growth and drug-induced death responses in gliomaspheres. Results in (a), (b), (c), (j), (k) and (l) show individual datapoints from three separate experiments. Data in (d) and (f) are mean ± s.d. from three independent experiments. Blots in (c), (k), and (l) are representative of three independent experiments. Asterisks in (h) indicate significance values from unpaired parametric t-tests comparing the proportion of cells assigned to be in S-phase in BSA vs ligand-treated spheroids. Asterisks in (j) indicate significance values from unpaired parametric t-tests comparing treated spheroids with their corresponding BSA+DMSO controls. For both, P ≤ 0.05, **P ≤ 0.01, ***P ≤ 0.001. Except for (i), all ligand treatments were 100 ng/mL. Scale bars = 400 µm.

We reasoned that DNA damaging agents like TMZ and VAL-083, and potentially kinase inhibitors like JCN037 and buparlisib, could require cell cycle progression to cause an accumulation of toxic lesions that lead to cell death in gliomaspheres independent of the canonical apoptosis pathway. Of note, of the 50 ligands we profiled for effects on cell death, only BMP2, BMP4, and OSM markedly reduced spheroid growth over time (**Fig. 4e,f**). Because these same ligands also protected against TMZ, VAL-083, JCN037 and buparlisib, we hypothesized that BMP2/4 blunted the killing of these agents by arresting cell cycle progression. Consistent with this hypothesis, genes involved in DNA replication and cell cycle progression were significantly downregulated in BMP2- and BMP4-treated GS025^N^ gliomaspheres as determined using RNA-sequencing (**Extended Data Fig. 7a**). Moreover, BMP2 and BMP4, but not BMP3, reduced the fraction of cells in S phase in both GS025^N^ and GS270^N^ gliomaspheres, as determined by flow cytometry (**Fig. 4g,h** and **Extended Data Fig. 7b**).

Previous studies suggested that BMP4 could upregulate the cyclin-dependent kinase inhibitors *CDKN1A* (p21) and *CDKN1B* (p27) to arrest the GBM cell cycle^57–59^. While BMP2/4 increased p21 and (less consistently) p27 abundance in a LDN193189-sensitive manner in GS025^N^, GS270^N^ and GS304^N^ gliomaspheres, rCRISPR-mediated disruption of *CDKN1A* or *CDKN1B* alone did not prevent BMP2 from modulating gliomasphere cell death or growth (**Extended Data Fig. 7c-g**). Given this potential functional redundancy, and test whether cell cycle arrest was sufficient to modulate gliomasphere cell death, we used the cyclin-dependent kinase 4/6 (CDK4/6) inhibitor palbociclib to block cell cycle progression at the level of retinoblastoma (Rb) protein function, which acts downstream of p21 and p27 (ref.^60, 61^). Palbociclib pre-treatment (1 µM, 24 h) arrested cell cycle progression (**Fig. 4g,h**) and reduced the lethality of TMZ, VAL-083, JCN037 and buparlisib, phenocopying the effects of BMP2 and BMP4 on compound-induced death (**Fig. 4i,j**). The effects of palbociclib were not additive with those of BMP2, indicating activity through a common mechanism. Palbociclib appeared likely to be acting in an on-target manner as GS270^N^ gliomaspheres, which harbor a loss-of-function mutation in *RB1*, were resistant to the effects of palbociclib on cell death (**Fig. 4i,j, Extended Data Fig. 7h**). The DNA damaging agents^3, 4^ TMZ and VAL-083 both increased γ-H2AX signals in GS025^N^ gliomaspheres and this effect was attenuated by co-treatment with either BMP2 or palbociclib (**Fig. 4k,l**). Together, these results suggest that BMP2/4 protect gliomaspheres from the lethal effects of DNA alkylating agents, and possibly some kinase inhibitors, by slowing cell cycle progression and thereby reducing proliferation-dependent cytotoxicity (**Fig. 4m**).

### BMP2/4 signaling primes gliomaspheres for lipid-dependent necrosis

Our results suggested that BMP2/4 had the unique ability to swtich cell death sensitivity in two directions at once, rendering gliomaspheres less sensitive to TMZ, VAL-083, JCN037 and buparlisib and simultaneously more sensitive to CIL56. We next sought to better understand the nature of this ligand-dependent cell death sensitivity switch. CIL56 triggers LiDN, a palmitate-dependent non-apoptotic cell death mechanism^24, 25^. LiDN can also be induced by tegavivint, a CIL56 analog that has advanced into clinical trials for lung, bone and blood cancers^25^. To our knowledge, tegavivint has not been tested in GBM. Across multiple patient-derived gliomasphere lines, BMP2 pretreatment increased sensitivity to both CIL56 and tegavivint as measured by HADES, and in each case, cell death was suppressed by the LiDN inhibitor TOFA^24, 25^ (**Fig. 5a,b**).

**Figure 5.**
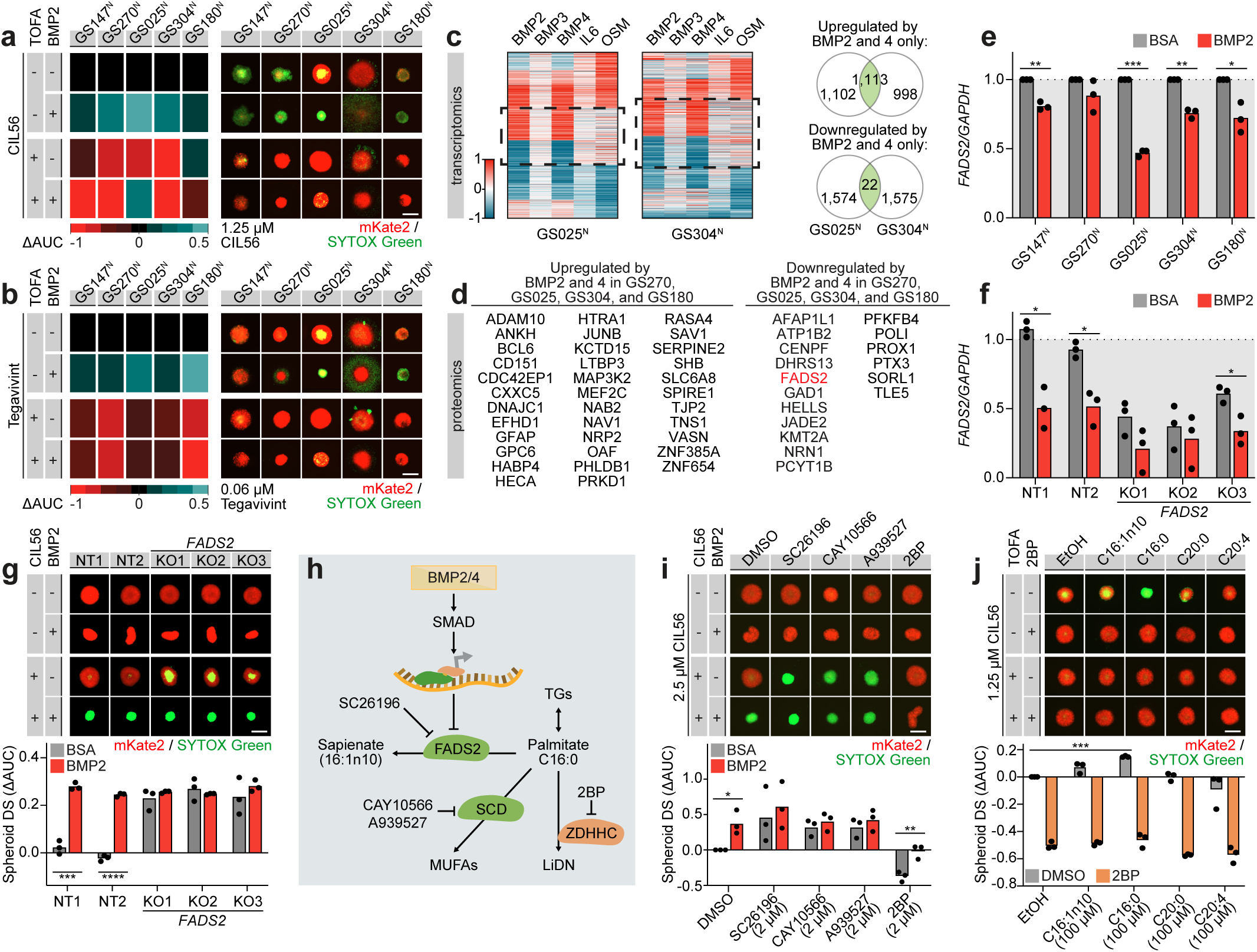
BMP signaling increases LiDN priming via FADS2. a, Effect of BMP2 on CIL56-induced death in gliomaspheres as quantified by HADES. CIL56 was tested across a 10-point, 2-fold dilution series starting at 10 µM. Heatmaps show changes in gliomasphere death ± BMP2 (100 ng/mL). ΔAUC = AUC_KO_ - average AUC_NT+BSA_. Representative images correspond to 1.25 µM CIL56. b, Effect of BMP2 on tegavivint-induced death in gliomaspheres as quantified by HADES. Tegavivint was tested across a 10-point, 2-fold dilution series starting at 1 µM. Heatmaps show changes in gliomasphere death ± BMP2 (100 ng/mL). ΔAUC = AUC_KO_ - average AUC_NT+BSA_. Representative images correspond to 60 nM tegavivint. c, RNA sequencing analysis for GS025^N^ and GS304^N^ gliomaspheres treated with the indicated ligands (100 ng/mL) relative to the BSA controls. Heatmaps were generated by unsupervised K-means clustering. Dotted black lines highlight the clusters of genes that are up- or down-regulated in response to BMP2 and BMP4 but not control ligands BMP3, IL6 and OSM. d, LC-MS-based proteomics analysis in the indicated patient-derived lines and conditions. Listed proteins are those that were consistently altered in the four patient-derived lines indicated and whose genes were identified in panel c as altered by BMP2 and BMP4 but not by control ligands BMP3, IL6 and OSM. e, Relative *FADS2* mRNA abundance measured by reverse transcription and quantitative PCR (RT-qPCR). f, Relative *FADS2* mRNA abundance measured by RT-qPCR in non-targeting control (NT) and *FADS2* knockout (KO) GS025^R^ gliomaspheres. g, Spheroid death in NT and FADS2 KO GS025^R^ gliomaspheres as determined by HADES. ΔAUC = AUC_KO_ - average AUC_NT+BSA_. h, Schematic of lipid-metabolic pathways influencing palmitate levels and LiDN sensitivity. i, Effect of pharmacological inhibitors indicated in panel h on GS025^N^ gliomasphere death response to CIL56. Representative images correspond to 2.5 µM CIL56. ΔAUC = AUC_treatment_ - AUC_BSA+DMSO_. j, Effect of exogenous lipids indicated in panel h on GS025^N^ gliomasphere death response to CIL56. Representative images correspond to 1.25 µM CIL56. ΔAUC = AUC_treatment_ - AUC_DMSO+EtOH_. Heatmaps in (a) and (b) summarize compound dose-response experiments from three independent experiments. Individual data points in (e), (f), (g), (i), and (j) are from three independent experiments. Asterisks in (e), (f), (g) and (i) indicate significance values from unpaired parametric t-tests comparing treated spheroids with their corresponding BSA+DMSO controls. Asterisks in (j) compare spheroids treated with exogenous lipids with their corresponding ethanol (EtOH) solvent control (*P ≤ 0.05, **P ≤ 0.01, ***P ≤ 0.001). Images in (a), (b), (g), (i) and (j) are representative of three independent experiments. All AUC values were obtained from 10-point two-fold dose-response curves (CIL56 = 10 µM – 0.02 µM; Tegavivint = 1 µM – 0.002 µM). All ligand treatments = 100 ng/mL. Scale bars = 400 µm.

How BMP2 and BMP4 primed gliomaspheres for LiDN was unclear. Palbociclib treatment did not alter gliomasphere sensitivity to CIL56 with or without concomitant BMP2 treatment, suggesting that cell cycle arrest did not explain LiDN priming (**Fig. 4i,j**). Building on the knowledge that LiDN priming required BMPR-SMAD signaling (**Fig. 3c,h**), RNA sequencing was used to identify 1,113 upregulated and 22 downregulated transcripts (FDR q < 0.05) shared between GS025^N^ and GS304^N^ gliomaspheres treated with BMP2 or BMP4, but not with control ligands that did not affect LiDN sensitivity (OSM, BMP3 or IL6) (**Fig. 5c**). Focusing on this gene expression space, liquid chromatography-mass spectrometry (LC-MS)-based proteomic analysis was used to identify 35 proteins that were significantly increased and 17 proteins that were significantly decreased by BMP2/4 across four gliomasphere lines (FDR q < 0.05) (**Fig. 5d**). Among the proteins decreased in abundance by BMP2/4 was the lipid metabolic enzyme fatty acid desaturase 2 (FADS2) (**Fig. 5d**).

Among other reactions, FADS2 catalyzes the conversion of palmitate (C16:0) to sapienate (C16:1)^62, 63^, a reaction that would limit the availability of palmitate for LiDN. We therefore hypothesized that BMP2/4 signaling increased primed gliomaspheres for LiDN by repressing FADS2 expression and increasing palmitate abundance. Consistent with this hypothesis, BMP2 decreased *FADS2* mRNA expression in multiple gliomasphere models, and this effect was blocked by co-treatment with LDN193189 or *SMAD4*^KO^ (**Fig. 5e** and **Extended Data Fig. 8a,b**). Treatment with the small molecule FADS2 inhibitor SC-26196 (ref.^64^) or rCRISPR-mediated *FADS2* disruption were both sufficient to sensitize GS025^N^ gliomaspheres to CIL56-induced death in the absence of BMP2/4 treatment (**Extended Data Fig. 8c** and **Fig. 5f,g**). Similar effects were observed in GS147^N^, GS270^N^ and GS304^N^ gliomaspheres treated with SC-26196, and the sensitizing effects of FADS2 inhibition were not additive with BMP2, suggesting activity through a shared pathway (**Extended Data Fig. 8d,e**). *FADS2* gene disruption did not alter gliomasphere sensitivity to TMZ, VAL-083, JCN037 or buparlisib, indicating that FADS2 specifically regulated LiDN priming (**Extended Data Fig. 8f**). By LC-MS-based lipidomic analysis we identified eighty-five lipid species increased by both BMP2 treatment and *FADS2* gene-disruption, most of which were triglycerides (**Extended Data Fig. 9a,b**). These triglycerides in BMP2-treated or *FADS2* gene-disrupted cells were enriched with saturated or monounsaturated 16- and 18-carbon acyl chains (**Extended Data Fig. 9c,d**), consistent with decreased flux to sapienate and ‘spillover’ of palmitate into triglycerides^65^.

Our results suggested that FADS2 downregulation upon BMP2/4 treatment increased LiDN sensitivity by increasing the pool of palmitate available for lethal palmitoylation (**Fig. 5h**)^24, 25^. In further support of this model, pharmacological inhibition of a distinct palmitate-consuming enzyme, stearoyl-CoA desaturase (SCD), using the small molecule inhibitors CAY10566 or A939527 increased 16:0-containing triglyceride species and was sufficient to sensitize GS025^N^ gliomaspheres towards CIL56, phenocopying *FADS2* gene disruption (**Fig. 5i, Extended Data Fig. 9e,f**). Conversely, the palmitoylation inhibitor 2-bromopalmitate (2-BP) protected cells from CIL56-induced death. Direct addition of exogenous palmitate similarly potentiated CIL56 lethality in GS025^N^ gliomaspheres, and this death was blocked by 2-BP (**Fig. 5j**). Together, these findings suggested that BMP2/4-SMAD signaling primed gliomaspheres for LiDN by suppressing FADS2 expression and increasing the pool of palmitate available for lethal protein palmitoylation.

### BMP-dependent cell death rewiring in DIPG

Our results thus far suggested that BMP2/4 could rewire GBM cell death sensitivity, reducing sensitivity to some agents and enhancing sensitivity to others. Diffuse intrinsic pontine glioma (DIPG) is a rare and highly lethal pediatric brain tumor, in which approximately 25% of patients carry activating mutations in the type I BMP receptor *ACVR1*^26–29^. Although ACVR1 is primarily activated by other BMP ligands including BMP7, it activates the same downstream SMAD signaling pathway as BMP2/4^48, 53^. Building on our findings in GBM, we formulated two hypotheses. First, that BMP7 would rewire cell death sensitivity in *ACVR1* wildtype DIPG, reducing sensitivity to alkylating agents and kinase inhibitors while enhancing sensitivity to LiDN. Second, that activating mutations in *ACVR1* would constitutively engage this switch and render DIPG cells insensitive to ligand-dependent modulation of cell death. To test these hypotheses, we applied HADES to rare patient-derived DIPG spheroids. BMP7 pretreatment (100 ng/mL, 24 h) protected *ACVR1* wildtype DIPG-6 and DIPG-38 spheroids against TMZ and JCN037, and sensitized to CIL56 and tegavivint (**Fig. 6a,b**). In these cells, BMP7 induced SMAD1/5/8 phosphorylation and expression of Inhibitor of DNA Binding 1 (ID1), a downstream target gene of SMAD signaling (**Fig. 6c**). In DIPG-36 spheroids harboring an ACVR1^G328E^ activating mutation^27^, biochemical markers of SMAD pathway activity were basally upregulated and BMP7 treatment did not alter sensitivity to compound treatment (**Fig. 6a-c**). Finally, BMP7 treatment decreased *FADS2* expression in the *ACVR1* wildtype but not mutant lines and blocking FADS2 activity with SC-26196 sensitized *ACVR1* wildtype DIPG spheroids to CIL56 and tegavivint-induced death in the absence of BMP7 (**Fig. 6d,e**). Thus, a conserved BMP-FADS2 axis seemed to regulate cell death sensitivity in DIPG spheroids and suggested that activating mutations in *ACVR1* may prime cells for LiDN.

**Figure 6.**
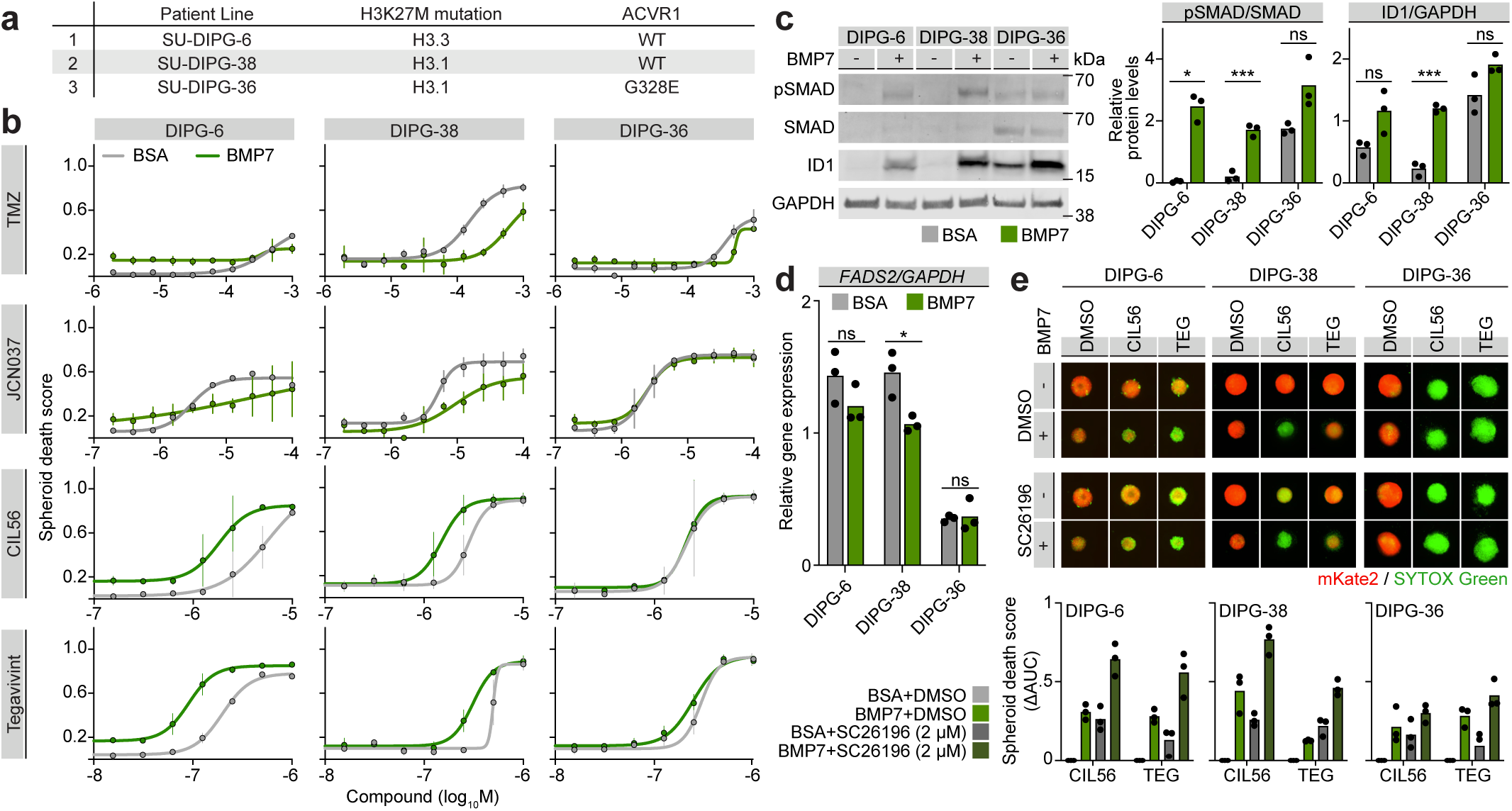
BMP signaling modulates death in DIPG patient-derived spheroids. a, Summary of patient-derived DIPG lines used in this study. b, Effect of BMP7 on DIPG spheroid death as determined by HADES. c, BMP pathway activation as measured by immunoblot of phosphorylated SMAD1/5/8 (pSMAD) and ID1, a downstream target gene of BMP signaling. Blots are representative of three independent experiments. d, Relative mRNA abundance determined using reverse transcription and quantitative PCR (RT-qPCR). e, Effect of the FADS2 inhibitor SC26196 (2 µM) on DIPG spheroid death induced by CIL56 or tegavivint. Bar graphs summarize ΔAUC (AUC_treated_-AUC_BSA+DMSO_) of compound dose-response experiments. Images are representative of three experiments and show spheroids treated with 2.5 µM CIL56 and 250 nM tegavivint. Scale bar = 400 µm. Data in (b) represent mean ± s.d. from three independent experiments. Individual data points in (c), (d), and (e) are from three independent experiments. Asterisks in (c) and (d) indicate significance values from unpaired parametric t-tests comparing BSA- and BMP7-treated spheroids (*P ≤ 0.05, **P ≤ 0.01, ***P ≤ 0.001). All ligand treatments = 100 ng/mL.

## DISCUSSION

HADES enables direct, high-throughput quantification of cell death in patient-derived GBM and DIPG spheroids, revealing phenotypes not captured in standard two-dimensional culture or by traditional viability-based assays that can only infer cell death. Using HADES, we found that BMP signaling potently regulates glioma cell death. Intriguingly, BMP2/4 in GBM and BMP7 in DIPG engaged a cell death switch that suppressed killing by DNA alkylating agents and kinase inhibitors while sensitizing cells to LiDN. We envision future applications of this HADES-based approach to study other molecules of interest present in the brain microenvironment (e.g., neurotransmitters), larger compound screens in BMP2/4-treated gliomaspheres, and extension of this technology to additional biological sample types.

Mechanistically, the BMP-dependent cell death switch operates through two separable arms. First, BMP2/4-induced proliferative arrest blunts the lethality of DNA alkylating agents and select kinase inhibitors. Decreased proliferation likely limits the accumulation of toxic DNA damage that drives cell death in response to alkylating agents like TMZ and VAL-083. How proliferative arrest reduces the lethality of kinase inhibitors in our models requires further elucidation, but we cannot exclude the possibility that these agents are only lethal in our models by acting on non-canonical targets. Second, BMP signaling reduced expression of the lipid desaturase FADS2, which likely sensitized cells to LiDN by increasing palmitate availability for lethal protein palmitoylation. Defining the critical palmitoylated substrates downstream of this pathway will be an important next step in understanding how lipid metabolism is coupled to LiDN execution in brain cancer.

Our findings reveal a therapeutic paradox in glioma. BMP signaling can restrain proliferation and is associated with less aggressive disease in patients^66, 67^ yet it simultaneously reduces sensitivity to the standard therapy, TMZ. However, this state may also expose a new vulnerability to LiDN. This may be particularly relevant in recurrent GBM, where BMP pathway gene expression is highly elevated, and in DIPG, where activating mutations in *ACVR1* occur in approximately 25% of tumors^26–29^. Recombinant human BMP4 has entered clinical testing for recurrent GBM^68^, raising the possibility of a “prime-then-kill” strategy in which BMP signaling is leveraged to sensitize tumors to LiDN-inducing agents. Although tegavivint is a clinical candidate LiDN inducer, its utility in brain cancer is uncertain and this compound may have poor blood-brain barrier penetration. Developing brain-penetrant LiDN-inducing agents could be one path towards engaging this new vulnerability in vivo.

## METHODS

### Patient-derived GBM gliomaspheres

All patient-derived GBM lines were obtained through the UCLA Institutional Review Board (IRB) protocol 10-000655 following explicit and informed consent from all patients. Patient age, sex, and mutational status are described in Extended Data Table 1. Gliomaspheres were established as previously described and maintained in gliomasphere media consisting of DMEM/F-12 (Corning, 10-092-CV), 1X B-27 supplement, minus Vitamin A (Thermo Fisher, 12587010), 1X P/S (Life Technologies, 15070-063), and 1X GlutaMax (Life Technologies, 35050-061), supplemented with heparin (Sigma, H3149, 5 μg/mL), EGF (Fisher, PHG0313, 50 ng/mL), and FGF-b (Fisher, PHG0263, 20 ng/mL). Cells were passaged every 5-7 days into fresh medium at a density of 75,000-100,000 cells/mL. For both routine passaging and experiment seeding, cells were pelleted, dissociated with TrypLE (Thermo Scientific, 12605028), filtered through a 40 or 70 µm cell strainer (Corning, 352340 or 352350), and counted with Trypan Blue (Sigma-Aldrich, T8154) using a Cellometer Auto T4 cell counter (Nexelcom). Gliomaspheres were used at fewer than 10 passages for the screen. For all other experiments excluding the overexpression, knockdown, and knockout studies, gliomaspheres were used at fewer than 25 passages. Cells were tested for *Mycoplasma* (Invivogen, rep-mysnc-50) and maintained in humidified tissue culture incubators (Thermo Scientific) at 37°C with 5% CO_2_.

### Patient-derived DIPG cells

DIPG lines were kind gifts from Dr. Michelle Monje (Stanford, California). Cells were maintained in tumour stem media consisting of Neurobasal-A Medium (Invitrogen, 10888-022), DMEM/F-12 (Corning, 10-092-CV), 1M HEPES Buffer Solution (Invitrogen, 15630-080), 1X MEM Sodium Pyruvate Solution (Invitrogen, 11360-070), 1X MEM Non-Essential Amino Acids Solution (Life Technologies, 11140-050), 1X GlutaMax (Life Technologies, 35050-061), 1X B-27 supplement, minus Vitamin A (Thermo Fisher, 12587010), and 1X P/S (Life Technologies, 15070-063), supplemented with 20 ng/mL EGF (Fisher Scientific, PHG0313), 20 ng/mL FGF-b (Fisher, PHG0263), 10 ng/mL PDGF-AA (Fisher Scientific, PHG0035), 10 ng/mL PDGF-BB (Fisher Scientific, PHG0045), and 2 µg/mL heparin (Sigma Aldrich, H3149). Cells were passaged every 6-10 days into 10 mL of conditioned media and 10 mL of 2X fresh media at a density of 75,000-100,000 cells/mL. For both routine passaging and experiment seeding, cells were pelleted, dissociated with TrypLE (Thermo Scientific, 12605028), filtered through a 70 µm cell strainer (Corning 352350), and counted using a Cellometer Auto T4 cell counter (Nexelcom). Cells were tested for *Mycoplasma* and maintained in humidified tissue culture incubators (Thermo Scientific) at 37°C with 5% CO_2_.

### Cancer cell lines

T98G^N^ cells were previously described^19^ and used at low passage numbers (P<10). Cells were grown in DMEM (Corning, 10-013-CV) supplemented with 10% heat inactivated FBS (Thermo Fisher Scientific, 26400044), and 1X P/S (Life Technologies, 15070-063). Cells were passaged every 2-3 days into fresh media at a density of 10,000 cells/cm^2^ using trypsin-EDTA (Thermo Scientific, 25200072). Cells were tested for *Mycoplasma* and maintained in humidified tissue culture incubators (Thermo Scientific) at 37°C with 5% CO_2_.

### Generation of nuclear mKate2-expressing glioma cells

GBM and DIPG patient-derived lines expressing nuclear localized mKate2 were generated by lentiviral spinfection with the Incucyte NucLight Red Lentivirus Reagent (Essen Bioscience, 4625) as follows. Spheroids were dissociated into single cell suspensions and plated in their respective media without P/S into 6 well plates at a density of 500,000 cells/mL, supplemented with polybrene (Sigma-Aldrich, H9268, 3.33 µg/mL) and lentivirus reagent. Plates were sealed with parafilm and spun at 800 x *g* for 90 min at 33°C. Immediately following spinfection, cells were pelleted and resuspended in fresh media. The following week, cells with similar levels of mKate2 fluorescence were sorted by fluorescence-activated cell sorting using a BD Aria II cytometer at the Stanford Shared FACS Facility. Cells were subsequently expanded, aliquoted, and frozen down.

### Quantification of gliomasphere growth and death using HADES

Gliomaspheres and DIPG spheroids stably expressing mKate2 were seeded into ultra-low attachment U-bottom 384 well plates (S-Bio, MS-9384UZ) at a density of 2,000 cells in 20 µL of medium per well. Plates were undisturbed for 24 h to allow cells to form spheroids, then treated with 20 µL of 2X ligands, then 20 µL of 3X compounds supplemented with 20 µL of 60 nM SYTOX Green (Life Technologies, S7020) for a final concentration of 20 nM SYTOX Green. Live cell images were acquired with an IncuCyte S3 (Essen Biosciences, 4647) using the following parameters: scan type spheroid, 10x objective in phase contrast, brightfield, green fluorescence (400 ms), red fluorescence (800 ms), and a spectral unmixing of 12% red contributing to green. Images were analyzed using the Incucyte software (Essen Biosciences, v2024A or v2024B). The analysis definition was as follows: brightfield sensitivity 50, adjust size 5 pixels, filters 3e4 µm^2^ minimum and 4e5 µm^2^ maximum, green segmentation adaptive with a threshold adjustment of 4 GCU and edge split off, filters 1000 µm^2^ minimum and mean intensity of 2.5, red segmentation adaptive with a threshold adjustment of 4 RCU and edge split off, filters 1000 µm^2^ minimum and mean intensity of 2.5. The integrated green and red fluorescence intensity scores, as well as the largest red area were exported to Excel for further data processing. Spheroid growth was calculated by normalizing the largest red area at *t=n* to the red area at *t=0*. For spheroid death, the red and green intensity scores were first normalized using the min-max scaling method in equation 1 then calculated into the spheroid death score using equation 2.

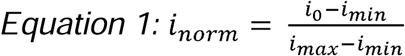

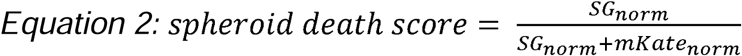

### Quantification of gliomasphere viability with CellTiter-Glo 3D

Gliomaspheres were seeded into ultra-low attachment U-bottom 384 well plates as described above. At the experiment endpoint, plates and CellTiter-Glo 3D viability assay reagent (Promega, G9681) were equilibrated to room temperature for 20 min. From each 384 well, originally containing 60 µL of culture media, 30 µL of medium was removed carefully, leaving the spheroid undisturbed. 30 µL of the CellTiter-Glo reagent was then added to each well. The plates were covered in foil and shaken for 5 min on an orbital shaker to induce cell lysis, then incubated at RT for 25m. Luminescence was recorded using a Cytation3 multimode plate reader (BioTek) with the following settings: 135 gain, 1 s integration time. The average background luminescence (measured from wells without cells and only containing the assay reagent and culture media) was subtracted from each experimental value, then normalized to the negative control of each treatment condition.

### Quantification of gliomasphere death with trypan blue

Cells were plated in 12 well plates at a density of 500,000 cells/mL. Five days following compound treatment, cells were pelleted, dissociated with TrypLE (Thermo Scientific, 12605028) and filtered through a 70 µm cell strainer (pluriSelect, 43-57070-51). Resulting cell suspensions were diluted at a 1:1 ratio with Trypan Blue (Sigma-Aldrich, T8154) and counted using a Cellometer Auto T4 cell counter (Nexelcom).

### Quantification of cell death and proliferation in adherent cells

Cell death in adherent T98G^N^ cells was quantified by scalable time-lapse analysis of cell death kinetics (STACK), as described previously^19, 69^. Live cell images were acquired with an IncuCyte S3 (Essen Biosciences, 4647) using the following parameters: scan type standard, 10x objective in phase contrast, green fluorescence (400 ms), red fluorescence (800 ms), and a spectral unmixing of 12% red contributing to green. Images were analyzed using the Incucyte software (Essen Biosciences, v2024A) with the following analysis definition: phase hole fill of 50 µm^2^ with a filter minimum of 300 µm^2^, green segmentation adaptive with a threshold adjustment of 10 GCU, edge split on with an edge sensitivity of -30 and a filter minimum of 40 µm^2^, red segmentation adaptive with a threshold adjustment of 1 RCU, edge split on with an edge sensitivity of -30, hole fill cleanup of 100 µm^2^ and filter minimum of 50 µm^2^, green and red filter minimum of 50 µm^2^. The number of green, red, and green+red (overlap) objects were exported to Excel for further data processing. Lethal fraction scores were calculated using the STACK method described in equation 3. Cell proliferation was calculated by normalizing the number of red objects at *t=n* by the number of red objects at *t=0*.

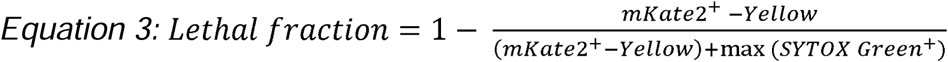

### Receptor-ligand analysis

Published RNA sequencing data for the GS025 gliomasphere line^15^ was used to identify cell surface receptors that were expressed with TPM > 0.5, then matched with their cognate ligands using CellChat^43^, ICELLNET^44^ and NATMI^45^. To prioritize clinically relevant receptor-ligand pairs, GBM patient survival probabilities (survival Z-score) for each receptor and ligand were obtained from The Cancer Genome Atlas (TCGA) program. Receptor-ligand pairs were ranked by the absolute value sum of their survival Z-scores (e.g., n=|Z_receptor_ + Z_ligand_|), as listed in Extended Data Figure 2b. Data cleaning, processing, and visualization were performed using Python v3.7.

### Ligand cell death modulation screen

Protein ligands were purchased (Extended Data Fig. 2b) and resuspended to 10,000 ng/mL with 0.1% BSA in PBS (Gemini Bio-Products, 700-100P), aliquoted, and frozen down to -80°C. Low passage GS025^N^ cells (P<10) were dissociated and seeded into ultra-low attachment U-bottom 384 well plates (S-Bio, MS-9384UZ) at a density of 2,000 cells in 20 µL of medium per well. 48 h post seeding, 20 µL of gliomasphere medium containing 200 ng/mL ligands was added to each well, resulting in a 1X ligand concentration of 100 ng/mL. The following day, 20 µL of gliomasphere medium containing SYTOX Green (Thermo Fisher Scientific, S7020, 20 nM) and lethal compounds (all tested in a 10-point, 2-fold dose-response series) was added at a 3X concentration, resulting in a final 1X concentration. Images and spheroid death scores were acquired every 24 h for 5 d, as described above. Wells with obvious technical errors were excluded from the analysis (e.g., wells with no spheroids, with debris, unfocused image acquisition). Five ligands were screened per week. A BSA control was included every week to control for batch-to-batch variations. The area under the curve of the spheroid death scores were calculated using Prism 10 (GraphPad), then subtracted by their corresponding lethal compound AUC values from the appropriate BSA controls. The Z-scores of each ligand were obtained for each ligand, relative to each lethal compound. Screen data were visualized using Morpheus (https://software.broadinstitute.org/morpheus).

### Protein extraction and immunoblotting

Cells were plated in 12 well plates at a density of 500,000 cells/mL. To harvest, spheroids were collected on ice, pelleted by centrifugation at 1000 x *g* for 5 min at 4°C, and washed with ice cold HBSS (Fisher Scientific, 14-175-079). Cells were then lysed with RIPA buffer containing p8340 (1:200, Sigma-Aldrich, P8340) and 5 mM NaF (Millipore Sigma, S6776), sonicated at 60% amplitude for 10 cycles of 1 s on and 1 s off using a Sonic Dismembrator (Cat# FB120110, Fisher Scientific, Hampton, NH), then cleared by centrifugation at 18,213 x *g* for 20 min at 4°C. Supernatants were collected and protein concentrations were quantified using the Pierce BCA Assay (Thermo Scientific, 23209). Standardized protein samples were mixed with 4X Bolt LDS Sample Buffer (Life Technologies, B0007) and 10X Bolt Sample Reducing Agent (Life Technologies, B0009) and heated at 70°C for 10 min. Proteins were separated by SDS PAGE on 10-, 12-, or 15-well Bolt 4-12% Bis-Tris Plus Gels (Thermo Fisher Scientific, NW04120BOX, NW04122BOX, or NW04125BOX) with the Duo Chameleon pre-stained protein ladder (LI-COR Biosciences, 928-60000) and dry transferred to nitrocellulose membranes using the iBlot2 or iBlot3 gel transfer devices (Fisher Scientific). Membranes were incubated with Intercept blocking buffer (LI-COR Biosciences, 927-70001) for 1 h at RT then with primary antibodies overnight at 4°C. The following primary antibodies and dilutions were used: rabbit anti-pSMAD1/5/8 (Cell Signaling Technology, 13820, 1:1500), rabbit anti-SMAD1 (Cell Signaling Technology, 6944, 1:1500), rabbit anti-SMAD4 (Cell Signaling Technology, 38454, 1:1000), rabbit anti-p21 (Santa Cruz, Sc-53870, 1:1000), rabbit anti-p27 (Cell Signaling Technology, 3686T, 1:1000), mouse anti-Rb (BD Pharmingen, 554136, 1:1000), mouse anti-Tubulin (Thermo Scientific, MS-581-P1, 1:10000), and mouse anti-GAPDH (Proteintech, 60004-1-Ig, 1:10000). The following day, membranes were washed thrice with Tris-buffered saline with Tween (TBST) for 5 min, incubated with secondary antibodies (1:10,000, LI-COR Biosciences, 926-68072 and 926-32213) for 1 h at RT, then washed thrice with TBST for 5 min. Blots were imaged using the Odyssey CLx or Odyssey M LI-COR imaging systems (LI-COR Biosciences). Band intensities were quantified using Fiji v2.14.

### RNA extraction and quantitative RT-PCR

Cells were seeded into 12 well plates at a density of 500,000 cells/mL. Spheroids were harvested on ice, pelleted by centrifugation at 1000 x *g* for 5 min at 4°C, and washed twice with ice cold HBSS (Fisher Scientific, 14-175-079). RNA was extracted using the RNeasy Plus Mini Kit (Qiagen, 74134) and cDNA synthesis was performed using the Applied Biosystems MultiScribe Reverse Transcriptase (Thermo Fisher Scientific, 43-112-35) with both the Applied Biosystems Oligo d(T) and Random Hexamer primers (Thermo Fisher Scientific, N8080128 and N8080127). Quantitative RT-PCR was performed using the Applied Biosystems SYBR Green PCR Master Mix (Thermo Scientific, 4309155) and an Applied Biosystems QuantStudio 3 instrument (Thermo Fisher Scientific). Relative expression levels were calculated using the ΔΔCT method. The following primers were used: *Actin* (forward: 5’-CATGTACGTTGCTATCCAGGC-3’, reverse: 5’-CTCCTTAATGTCACGCACGAT-3’), *BMPR1A* (forward: 5’-CCAAGGAAAGCCCGCAATTG-3’, reverse: 5’-CACCCTGGTATTCAAGGGCAC-3’), *FADS2* (forward: 5’-CTCATCACGGCCTTTGTCCT-3’, reverse: 5’-TGGAAGATGTTAGGCTTGGCG-3’).

### Cloning

For *SMAD4* overexpression experiments, a plasmid containing the *SMAD4* cDNA was purchased (Addgene, Plasmid#14859) then subcloned into pLenti CMV Puro DEST (Addgene, Plasmid# 17452) by Gibson assembly (New England Biolabs, E2611S) using the following primers: *SMAD4* (forward: 5’-aggtctatataagcagagctATGGACTACAAGGACGAC-3’, reverse: 5’-TCAGTCTAAAGGTTGTGG-3’), pCMV Lenti (forward: 5’-acccacaacctttagactgaAATCAACCTCTGGATTACAAAATTTG-3’, reverse: 5’-AGCTCTGCTTATATAGACC-3’). A SMAD4^I500V^ dominant negative mutant^54^ was generated by site directed mutagenesis (New England Biolabs, E0554S0) of the pLenti-SMAD4 plasmid using the following forward and reverse primers: 5’-CCCAAGACAGAGCGTC-3’, and 5’-TAATCCGGTCCCCAG-3’. Cas9 expressing gliomaspheres were generated by introducing a plasmid containing Cas9 and mKate2 into low passage (P<5) GS025 cells. This plasmid was engineered by Gibson assembly of the Cas9 cassette and vector backbone (Addgene, Plasmid# 52962) with an mKate2 fragment generated by Twist Biosciences. The following forward and reverse primers were used to amplify the Cas9 cassette and vector backbone with sequences to overlap the mKate2 fragment: 5’-gcaaactggggcacagatgaAATCAACCTCTGGATTACAAAATTTG-3’ and 5’-ttaatcagctcgctcaccatCGGTCCAGGATTCTCTTC-3’. All constructs were validated by Sanger sequencing, followed by whole plasmid sequencing (Elim Biopharm).

### Generation of gliomasphere overexpression and knockdown lines

Lentivirus particles used for the generation of overexpression and knockdown gliomasphere lines were produced using Lenti-X cells (Takara Bio). Lenti-X cells were plated in 6 cm dishes coated with 1X Poly-D-lysine (Thermo Scientific, A3890401). Immediately before transfection, the Lenti-X cells were gently washed with 1X PBS (VWR, VWRV0780) and replaced with gliomasphere media minus P/S. Overexpression or shRNA knockdown plasmids were transfected along with the lentiviral packaging plasmids psPAX2 and pMD2.G (Addgene, Plasmid #12260 and 12259) using the TransIT-Lenti transfection reagent (Mirus, MIR 6606) following the manufacturer’s protocol. 6-12 h post transfection, media was replaced with fresh media containing 1X ViralBoost reagent (Alstem Cell Advancements, VB100). Supernatants were collected 48 h later, spun down at 450 x *g* for 5 min, passed through a 0.45 µm filter, aliquoted, and frozen down. Cells were spinfected with the viral media as described above, then subjected to puromycin selection (Thermo Fisher Scientific, A11138-03, 1 µg/mL) 1 week post spinfection. shRNA knockdown lines were maintained in 0.5 µg/mL puromycin and used three passages or fewer post-spinfection. Plasmids used to generate these lines are described in Extended Data Table 3.

### Generation of gliomasphere knockout lines

GS025-Cas9 (GS025^R^) gliomaspheres were engineered as described above. Cells with similar levels of Cas9-mKate2 levels were sorted by fluorescence-activated cell sorting with a BD Aria II cytometer at the Stanford Shared FACS Facility. GS025^R^ cells were expanded, aliquoted, and frozen down. Single guide RNAs were purchased from Integrated DNA Technologies (Coralville, IA) resuspended in nuclease-free water and electroporated into GS025-Cas9 cells using the Human Stem Cell Nucleofector Kit 1 and Amaxa Nucleofector II electroporator (Lonza Bioscience) with program A033. 1 week post electroporation, knockouts were verified by immunoblot, then expanded, aliquoted, and frozen down. All knockout lines and sgRNA sequences are described in Extended Data Table 3.

### Cell cycle profiling

Cells were seeded into 24 well plates (Corning, 3764) at a density of 1 x 10^6^ cells/mL. The following day, cells were treated with ligands or compounds. Following a 24 h incubation period, cell pellets were collected, dissociated with TrypLE (Thermo Scientific, 12605028) for 5 min, then diluted with Ca/Mg^2+^ free HBSS (Fisher Scientific, 14-175,079). Pellets were washed twice with Ca/Mg^2+^ free HBSS, then fixed by dropwise addition of ice cold 70% ethanol while vortexing. The following day, fixed cells were equilibrated to RT and washed once with Ca/Mg^2+^ free HBSS. Cells were stained with a FITC conjugated Ki-67 antibody (Thermo Scientific, 11-5698-82, 0.25 µg) diluted in HBSS for 1 h at RT, then incubated with a solution containing propidium iodide (Abcam, ab14083, 50 µg/mL), RNase A (Qiagen, 19101, 100 µg/mL), MgCl_2_ (Thermo Fisher, AM9530G, 2 mM) and HBSS for 1 h at 37°C. Stained cells were pelleted and resuspended in Ca/Mg2+ free HBSS and passed through a 35 µm strainer (Corning, 352235). Data were collected using an Attune NxT Flow Cytometer (Thermo Fisher Scientific) and analyzed using the FCS Express Flow Cytometry Software (De Novo Software). Cell cycle status was assigned as described^70^.

### BH3 profiling

Cells were plated and incubated with ligands or compounds in 12 well plates (Corning, 3413). Following a 24 h incubation period, cells were dissociated with TrypLE, passed through a 70 µM filter, and counted with a Cellometer Auto T4 cell counter (Nexelcom). Cells were resuspended in MEB buffer (150 mM mannitol, 10 mM HEPES-KOH, 50 mM KCl, 0.02 mM EGTA, 0.02 mM EDTA, 0.1% BSA, and 5 mM succinate) at a density of 50,000 cells/50 µL then added to 96 well plates containing 50 µL of 0.002% digitonin plus a 2X titration of BIM peptides (0 µM, 0.01 µM, 0.03 µM, 0.1 µM, 0.3 µM, 1 µM, 3 µM, 10 µM). Plates were incubated at RT for 50 min, fixed with 4% paraformaldehyde (Santa Cruz Biotechnology, sc-281692) for 10m, then neutralized with N2 buffer (1.7 M Tris and 1.25 M glycine, pH 9.1) for 5 min. Samples were stained overnight with a 10X staining solution (10% BSA and 2% Tween20 in PBS) containing DAPI (Sigma-Aldrich, D9542, 1:100) and an Alexa Fluor 488 conjugated anti-cytochrome C antibody (BioLegend, 612308, 1:400). Cytochrome c release was quantified with an Attune NxT Flow Cytometer (Thermo Fisher Scientific) and the FCS Express Flow Cytometry Software (De Novo Software) and gated based on the positive and negative lethal controls (25 µM alamethicin and DMSO). To obtain the delta priming score, the area under the curves of the cytochrome c release in response to the titration of BIM peptides were calculated, then normalized to the negative ligand or compound controls (BSA or DMSO). All conditions were run with two biological replicates and at least three experimental replicates.

### Caspase-Glo assay

Cells were seeded into ultra-low attachment U-bottom 384 well plates as described above. On the day of harvest, Caspase-Glo reagent (Promega, G8091) was equilibrated to RT and prepared as instructed by the manufacturer. From each well originally containing 60 µL of culture media, 30 µL of medium was carefully removed and replaced with 20 µL of the Caspase-Glo reagent. Plates were covered in foil and incubated on an orbital shaker for 1 h at RT. Luminescence was recorded using a Cytation3 multimode plate reader (BioTek) with the following settings: 135 gain, 1 s integration time. Luminescence readings from each well were subtracted by the average blank value (measured from wells without cells and only containing the assay reagent and culture media) then normalized to the negative control for each treatment condition.

### Transcriptomics

Low passage (P<10) cells were seeded into 6 well plates (Corning, 3471) at a density of 500,000 cells in 2 mL of media. 48 h post seeding, ligands were added directly into the culture media at a concentration of 100 ng/mL. Spheroids were harvested and collected for RNA as described above. Isolated RNA was assessed for quality and concentration using an Agilent 2100 Bioanalyzer at the Stanford Protein and Nucleic Acid facility. Library generation, read sequencing, and data cleanup were performed by Novogene on an Illumina HiSeq 4000 platform. The log_2_FC scores were obtained by normalizing to the BSA control, then visualized and clustered by k-means clustering using Morpheus (https://software.broadinstitute.org/morpheus). All conditions consisted of two independent experimental replicates.

### Proteomics

Low passage (P<15) cells were seeded into 6 well plates at a density of 1 x 10^6^ cells in 2 mL of media and treated with ligands as described above. Gliomaspheres were collected on ice, pelleted, and washed with HBSS (Fisher Scientific, 14-175-079). Cell pellets were flash frozen with liquid nitrogen and stored at - 80°C. Cell pellets were lysed on ice with RIPA buffer (Boston BioProducts, BP-115) containing protease and phosphatase inhibitor cocktail (Thermofisher, 1861282) for 30 min. Protein concentration was measured using BCA (Thermo Scientific, 23225). 200 μg of protein was reduced and alkylated using 5 mM TCEP (Thermo Scientific, 77720) and 10 mM CAA (Thermo Scientific, A39720). Reduced and alkyated proteins were then bound to carboxylate-modified beads following an SP3 protocol using an automated KingFisher instrument, essentially as described^71^. A 10:1 bead to protein mass ratio was used for binding of proteins to beads, and Trypsin/Lys-C (Thermo Scientific, A41007) was used to digest proteins into peptides at a 1:50 enzyme to protein mass ratio. Peptides were eluted from SP3 beads using 2% DMSO, dried down and resuspended in 2% acetonitrile (Thermo Scientific, 51101) / 0.1% trifluoroacetic acid (Thermo Scientific, 28904) in LC-MS grade water, vortexed, and sonicated for 15 minutes at room temperature. Following a brief centrifugation, samples were placed on a magnetic separator (Invitrogen, 12321D) to remove residual beads, and 20 μL of the clarified peptide solution was transferred to 9 mm autosampler vials for UHPLC-MS/MS analysis.

Peptide samples were separated using a Vanquish Neo UHPLC system (Thermo Scientific, VN-S10-A-01). Peptides were loaded onto a 300 µm x 5 mm PepMap trap column) at 10 µL/min with 100% mobile phase A (0.1% formic acid in water) for 4 cycles at a maximum pressure of 800 bar. Following trapping, peptides were separated on a 150 μm x 15 cm analytical IonOpticks column maintained at 60°C using a binary gradient at a flow rate of 3.2 µL/min. Mobile phase A consisted of 0.1% formic acid in water, and mobile phase B consisted of 0.1% formic acid in 100% acetonitrile. The 15 min gradient profile included an initial equilibration at 2.7 µL/min, followed by peptide elution with increasing organic content. The autosampler was maintained at 4°C throughout the analysis. Tandem mass spectrometry was performed on an Orbitrap Astral mass spectrometer (Thermo Scientific, BRE725660) equipped with a nanospray ionization (NSI) source. The instrument was operated in positive ion mode with spray voltage of +1900 V. The ion transfer tube was set to 290°C. An EASY-IC internal calibrant was used for real-time mass calibration throughout the analysis. The instrument method employed a data-independent acquisition (DIA) strategy with a total cycle time of 15 min.

#### MS1 survey scans

Full-scan MS spectra were acquired in the Orbitrap mass analyzer over a range of 380-980 m/z with a resolution of 240,000 (at 200 m/z). The normalized automatic gain control (AGC) target was set to 500%, with a maximum injection time of 3 ms and an RF lens setting of 40%. Advanced peak determination was enabled with an expected peak width of 6 sec and a default charge state of +2.

#### DIA MS2 scans

Data-independent acquisition was performed using the Astral mass analyzer with 149 variable isolation windows spanning the precursor mass range of 380-980 m/z. Isolation windows were 4 m/z wide with 0 m/z overlap, and window placement optimization was enabled. Precursor ions were fragmented using higher-energy collisional dissociation (HCD) with a normalized collision energy of 25%. Fragment ions were detected in the Astral analyzer scanning from 150-2000 m/z with a maximum injection time of 7 ms, normalized AGC target of 500%, and RF lens of 40%. The DIA scan cycle time was 0.6 sec, with loop control set to time-based acquisition throughout the 15 min gradient.

Raw mass spectrometry data files were converted to mzML format (full peak profile) and processed using DIA-NN (Data-Independent Acquisition by Neural Networks) version 1.9.1 (ref.^72^). Peptide and protein identification were performed using a double-pass search strategy against the UniProt human reference proteome database (version: 15 August 2024 containing 20,406 protein isoforms) and utilized a deep learning-generated predicted spectral library containing 6,080,596 precursors across 2,561,169 elution groups. Trypsin/Lys-C was specified as the digestion enzyme with cleavage at lysine and arginine residues (K*/R*), allowing up to 2 missed cleavages. Carbamidomethylation of cysteine was set as a fixed modification, and N-terminal methionine excision was enabled. Mass accuracy optimization was performed automatically using the first run in the dataset. The search employed neural network-based algorithms for both spectral library generation and targeted peak selection. A library-free double-pass search was performed in which DIA-NN first generated a project-specific spectral library from the acquired raw DIA MS data, then reanalyzed all runs against this empirically derived library. Retention times were aligned empirically across all runs using the RT profiling algorithm. High-precision quantification mode was enabled for improved precision of precursor intensity measurements. Protein identification and grouping were performed using heuristic protein inference algorithms with relaxed settings to minimize redundancy in protein groups. Gene-level inference was performed using only proteotypic peptides (peptides mapping uniquely to a single gene).

Precursor and protein identifications were filtered to a false discovery rate (FDR) of 1% using target-decoy competition against a reversed protein database and neural network-based q-value estimation. Precursor-level q-values were calculated independently for each run, followed by protein-level q-value calculation using the picked protein FDR control strategy. Global protein group q-values were calculated across all runs to ensure consistent protein-level FDR control.

Label-free quantification was performed using the extracted ion chromatograms (XICs) with a 10 sec extraction window centered on the peak apex. Precursor quantities were integrated across all isotopic peaks and charge states. Protein quantities were calculated using the MaxLFQ algorithm, which performs intensity-based absolute quantification using shared peptides while minimizing the impact of missing values. Cross-run normalization was applied to correct for systematic differences in total signal intensity between samples. The final dataset comprised 10,192 protein groups passing the 1% global FDR threshold, representing 9,170 genes at the precursor level and 8,568 genes at the protein level using proteotypic peptide inference.

### Lipidomics

GS025^R^ non-targeting or gene targeted cells were seeded into 6 well plates (Corning, 3471) at a density of 2E6 cells in 2 mL of media. 48 h post seeding, inhibitors or ligands were added directly into the culture media. The following day, spheroids were spun down, dissociated with TrypLE (Thermo Scientific, 12605028) for 4 min, then resuspended diluted with Ca/Mg^2+^ free HBSS (Fisher Scientific, 14-175,079) and filtered through a 70 µm cell strainer (pluriSelect, 43-57070-51). Viable cells were counted with Trypan Blue (Sigma-Aldrich, T8154) using a Cellometer Auto T4 cell counter (Nexelcom), then pelleted.

A 1:1 (v/v) butanol/methanol solution containing 10 mM ammonium formate was added to each cell pellet at a ratio of 1E6 cells per 100 µL extraction solvent. Samples were vortexed and sonicated for 1 h at RT. Following sonication, samples were centrifuged at 15,000 rpm for 10 min at 20°C. Subsequently, 76 µL of the supernatant were transferred to glass vials with inserts and spiked with 4 µL of Avanti’s SPLASH II Lipidomix Mass Spec standard as internal standards. Dynamic Multiple reaction monitoring (dMRM) LC/MS analysis was performed using an Agilent 1290 Infinity II Bio LC system coupled to an Agilent 6495D Triple Quadrupole Mass Spectrometer. An aliquot of 3 µL from each extract was injected onto a ZORBAX Eclipse Plus C18 column (100 x 2.1 mm, 1.8 um), with the column temperature maintained at 45°C.

The mobile phases consisted of 10 mM ammonium formate and 5 µM deactivator additive in a 5:3:2 (v/v/v) mixture of water:acetonitrile:2-propanol and 10 mM ammonium formate in a 1:9:90 (v/v/v) mixture of water:acetonitrile:2-propanol. The chromatographic gradient was as follows: initial,15% B; 0-2.5 min, linear increase from 15% to 50% B; 2.5-2.6 min, ramp to 57% B; 2.6-9 min, linear increase from 57% to 70% B; 9.0-9.1 min, ramp to 93% B; 9.1-11 min, linear increase from 93% to 96% B; 11.0-11.1 min, ramp to 100% B; 11.1-14.0 min, hold at 100% B; 14-14.2 min, decrease from 100% to 15% B; and 14.2-18 min hold at 15% B for re-equilibration. For the 6495D Triple Quadrupole system, the following parameters were applied: polarity switching mode was used, with capillary voltages set to +3500 V for positive ion mode and -3000V for negative ion mode. The drying gas temperature was maintained at 150°C with a flow rate of 17 L/min, while the sheath gas temperature was set to 200°C with a flow rate of 10 L/min. The nozzle voltage was 1000 V for positive ion mode and -1500 V for negative ion mode. Collision energies were optimized for each lipid class. A total of 667 MRMs transitions were monitored (Positive: 648, negative: 19) with a cycle time of 750 ms.

The dMRM LC/MS raw data were processed using Agilent MassHunter Quantitative Analysis Software (Version 12.1). A predefined dMRM method, including all MRM transitions and retention times (RTs) for 762 targeted lipids and 14 internal standards, was used for peak extractions. Each lipid peak integration was manually inspected to ensure accurate and consistent quantitation. The processed data was then exported to an excel file.

### Analysis of published single cell and single nuclei RNAseq data

Single nuclei RNA sequencing data^49^ was obtained from GSE228500. This dataset comprises specimens from two epilepsy control patients, 10 patients with primary IDH1-wildtype high grade glioma and eight patients with recurrent IDH1-wildtype high grade glioma. We classified neoplastic versus non-neoplastic nuclei based on chromosomal copy number variation (CNV) inference. Cells that were annotated as transformed tumor cells were subsetted and marker genes were identified for each patient using the drop-out curve method, as described previously^49^. We took the union of the resulting marker sets to cluster and embed the merged 20-patient dataset. Merged snRNA-seq count data was loaded into R and normalized using the Seurat package.

Each tumor cell was scored using the package AUCell in R (version 4.5.1) for each of the six lineage gene sets described by Neftel et al^31^: Mesenchymal 1 (MES1), Mesenchymal 2 (MES2), Astrocyte-like (AC), NPC-like 1 (NPC1), NPC-like 2 (NPC2), OPC-like (OPC). For each transformed cell, the highest of the six lineage scores via AUCell defined its lineage classification. Each cell also underwent cell cycle scoring using the “CellCycleScoring” function in Seurat and the published proliferation gene sets G1/S and G2/M^31^, which classified each tumor cell into “G1/S” and “G2/M” groups. Tumor associated macrophages (TAMs) were classified into proliferative, monocyte-derived and microglia-derived TAMs. Non-neoplastic tumor associated astrocytes were classified into type 1 (protoplasmic), type 2 (reactive with expression of oligodendroglial and neuronal genes) and type 3 (reactive with expression of inflammatory genes)^49^. These classifications were then used to analyze expression of *BMP2*, its receptors (*BMPR1A*, *BMPR1B*, *BMPR2*) and its transcriptional targets (*ID1*) in different glioma cell states and cells in the glioma microenvironment using the DotPlot function in Seurat.

### Statistical analysis

Unless otherwise specified, all experiments consisted of at least three independent experimental replicates and all comparisons were made with two-tailed, unpaired Student’s t-test. Data were graphed and analyzed using Prism 10 (GraphPad) or Python. For all figures, data represent mean ± s.d. and p > 0.05 = ns; p < 0.05 = *; p < 0.01 = **; p < 0.001 = ***; p < 0.0001 = ****.

## DATA AVAILABILITY

All data needed to evaluate the conclusions stated in the paper are presented in the paper and/or the supplementary materials. Cell lines and plasmids can be requested to the corresponding author. RNA sequencing data from untreated GS025 gliomaspheres were published previously and deposited to dbGaP under the accession number phs003286. All other RNA sequencing data, generated in this study, were deposited to the GEO under the accession number GSE312023.

## ACKNOWLEDGEMENTS

We thank Michelle Monje for DIPG patient-derived lines, Yuqin Dai and the Stanford Metabolomics Core for help with lipidomics experiments, the Stanford Shared FACS Facility, and members of the Dixon and Nathanson labs for feedback on the manuscript. The research was supported by funding from the Charlie Teo Foundation (to D.A.N. and S.J.D.) and the National Institutes of Health (R01NS143119 to D.A.N. and S.J.D.).

## AUTHOR INFORMATION

### Contributions

W.C.L., J.S., D.A.N., and S.J.D. conceived and designed this study. W.C.L. performed most of the experiments. J.S. and D.C. performed proteomics on the gliomaspheres. A.G. assisted with some immunoblots. M.B. performed the single nuclei data analysis. W.C.L. and S.J.D. wrote the original draft of this paper. W.C.L., J.S., D.A.N., S.J.D. reviewed and edited the paper. S.J.D. and D.A.N. were responsible for funding acquisition.

## ETHICS DECLARATION

The authors declare no competing interests.

---

**Extended Data Figure 1.**
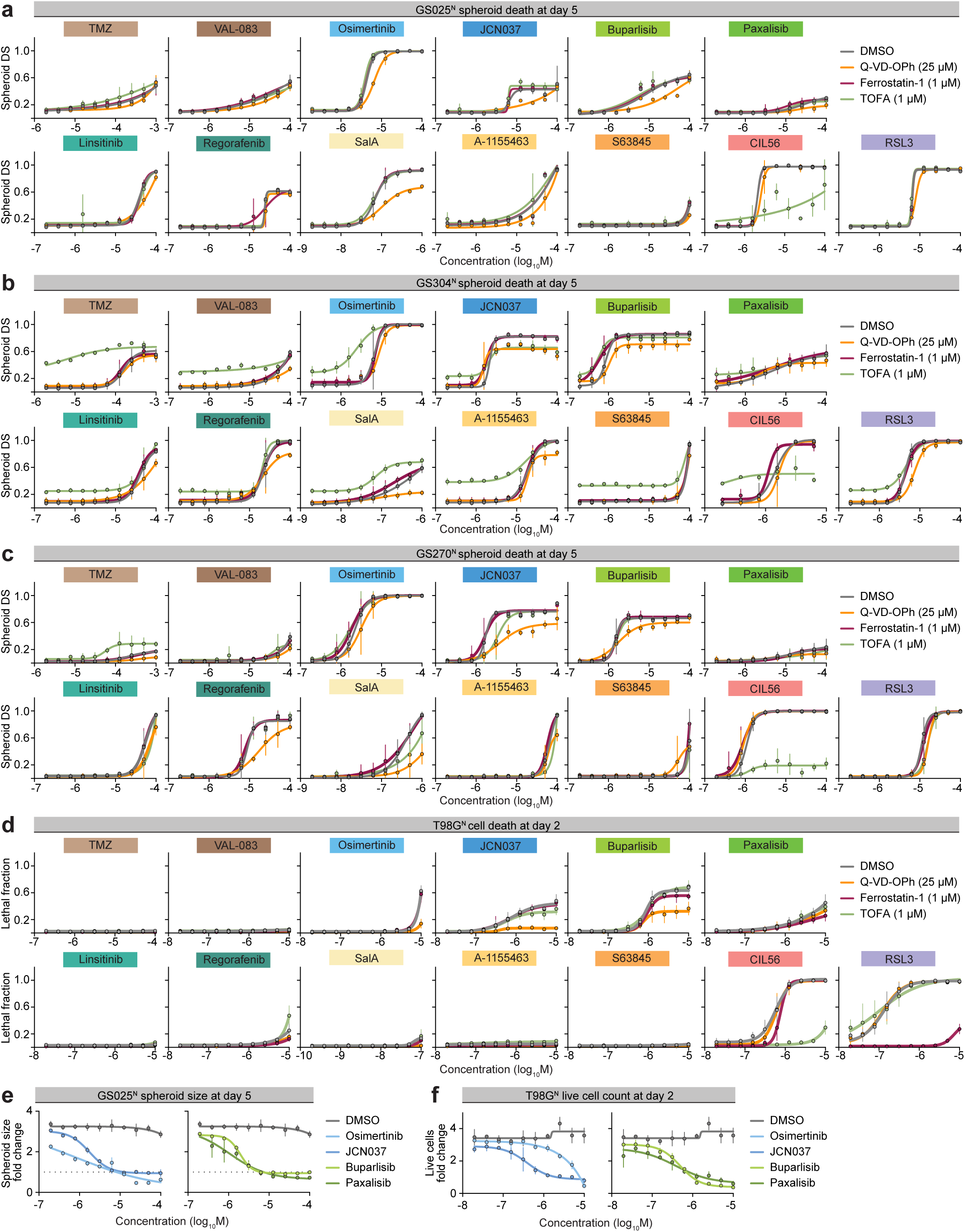
High-throughput analysis of gliomasphere cell death. a, Compound dose response curves in the GS025^N^ cells with and without apoptosis (Q-VD-OPh), ferroptosis (Ferrostatin-1), or LiDN (TOFA) inhibitors. b, Compound dose response curves in the GS304^N^ cells with and without apoptosis, ferroptosis, or LiDN inhibitors. c, Compound dose response curves in the GS270^N^ cells with and without apoptosis, ferroptosis, or LiDN inhibitors. d, Compound dose response curves in adherent T98G^N^ cells with and without apoptosis, ferroptosis, or LiDN inhibitors. e, Effect of kinase inhibitors targeting growth factor signaling pathways on GS025^N^ spheroid growth. f, Effect of kinase inhibitors targeting growth factor signaling pathways on T98G^N^ proliferation. All data represent mean ± s.d. from three independent experiments.

**Extended Data Figure 2.**
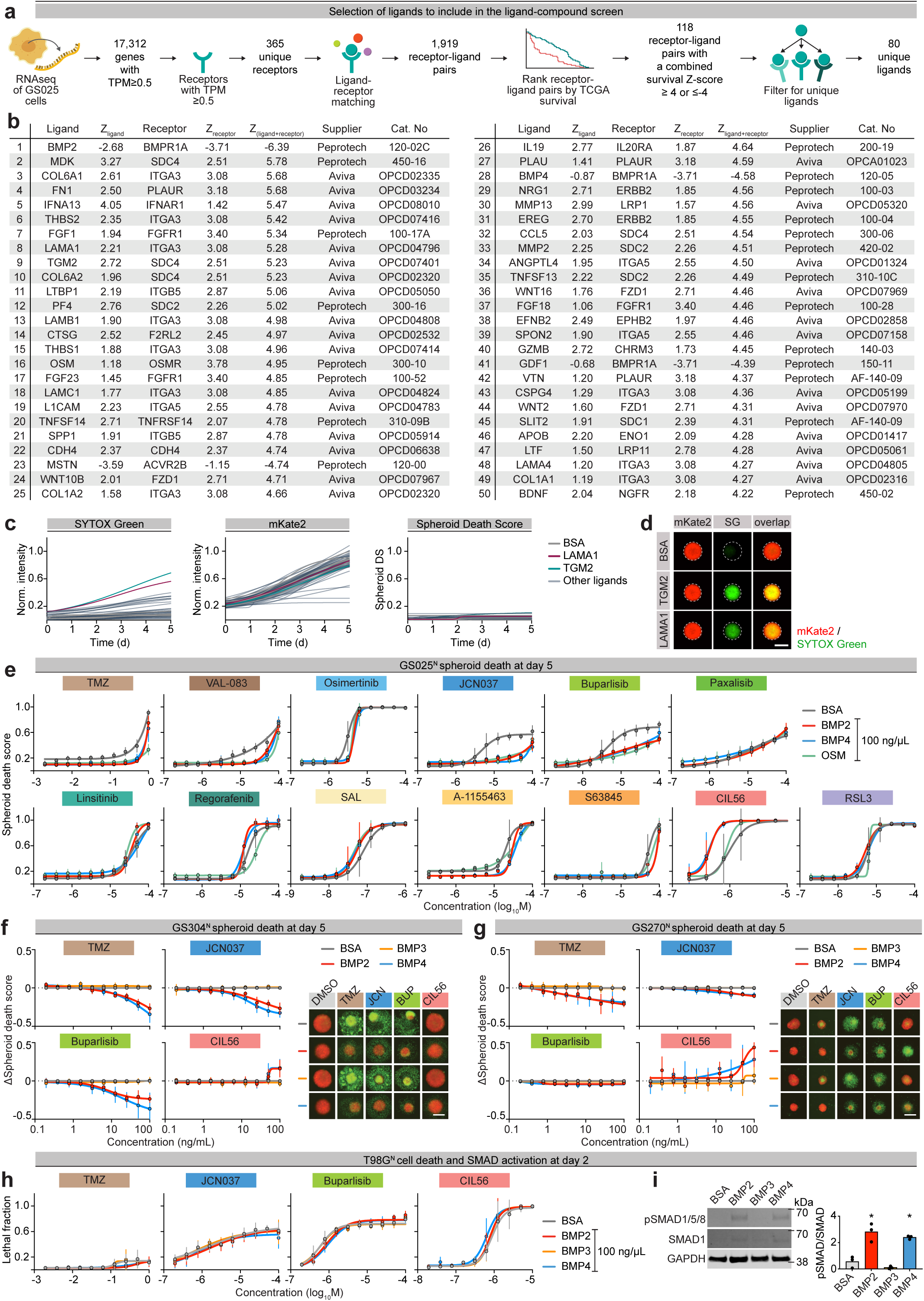
Selection and validation of ligands. a, Overview of the analysis used to nominate ligands for screening. b, List of the 50 ligands included in the screen with their corresponding receptors and TCGA survival Z-scores c, Baseline effects of ligands (100 ng/mL) on GS025^N^ gliomaspheres in the absence of lethal compounds, shown as normalized mKate2 and SYTOX Green intensities and derived spheroid death scores. d, Representative images highlighting ligand conditions (LAMA1 and TGM2) that increased SYTOX Green signal without a corresponding decrease in mKate2 or spheroid growth. e, Compound dose-response curves illustrating OSM-, BMP2-, and BMP4-mediated modulation of spheroid death (corresponding to bar graph summaries in Figure 1g). f, Ligand dose-response curves illustrating BMP2- and BMP4-mediated modulation of spheroid death in GS304^N^ gliomaspheres. g, Ligand dose-response curves illustrating BMP2- and BMP4-mediated modulation of spheroid death in GS270^N^ gliomaspheres. h, Ligand effects on T98G^N^ cell death response quantified by imaging. i, Immunoblot analysis of SMAD1/5/8 phosphorylation (pSMAD) following ligand treatment. Blots are representative of four independent experiments. Data in (e), (f), (g), and (h) represent mean ± s.d. from three independent experiments. Images in (f) and (g) are representative of three independent experiments. Scale bars = 400 µm.

**Extended Data Figure 3.**
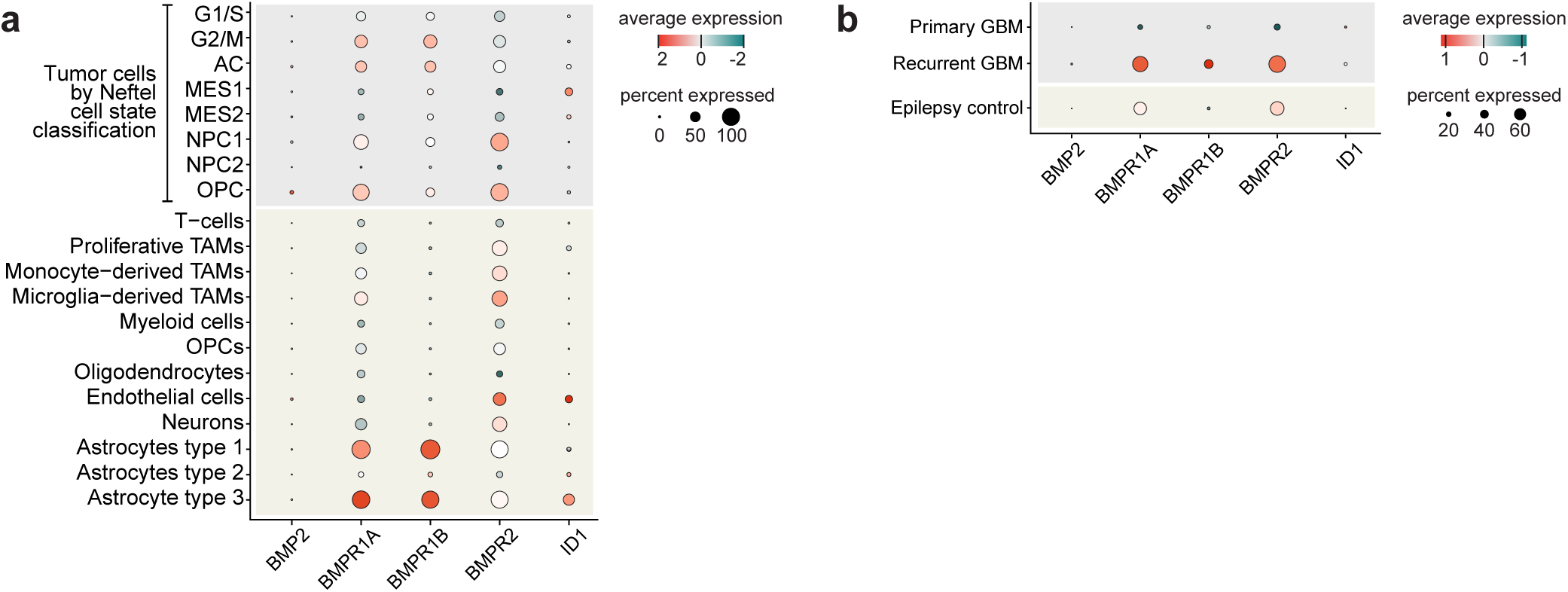
Effects of ligands on GBM cell states. a, Dot plots showing expression of BMP receptors across cell types in GBM patient snRNA-seq data. b, Expression of BMP receptors in primary and recurrent GBM tissue.

**Extended Data Figure 4.**
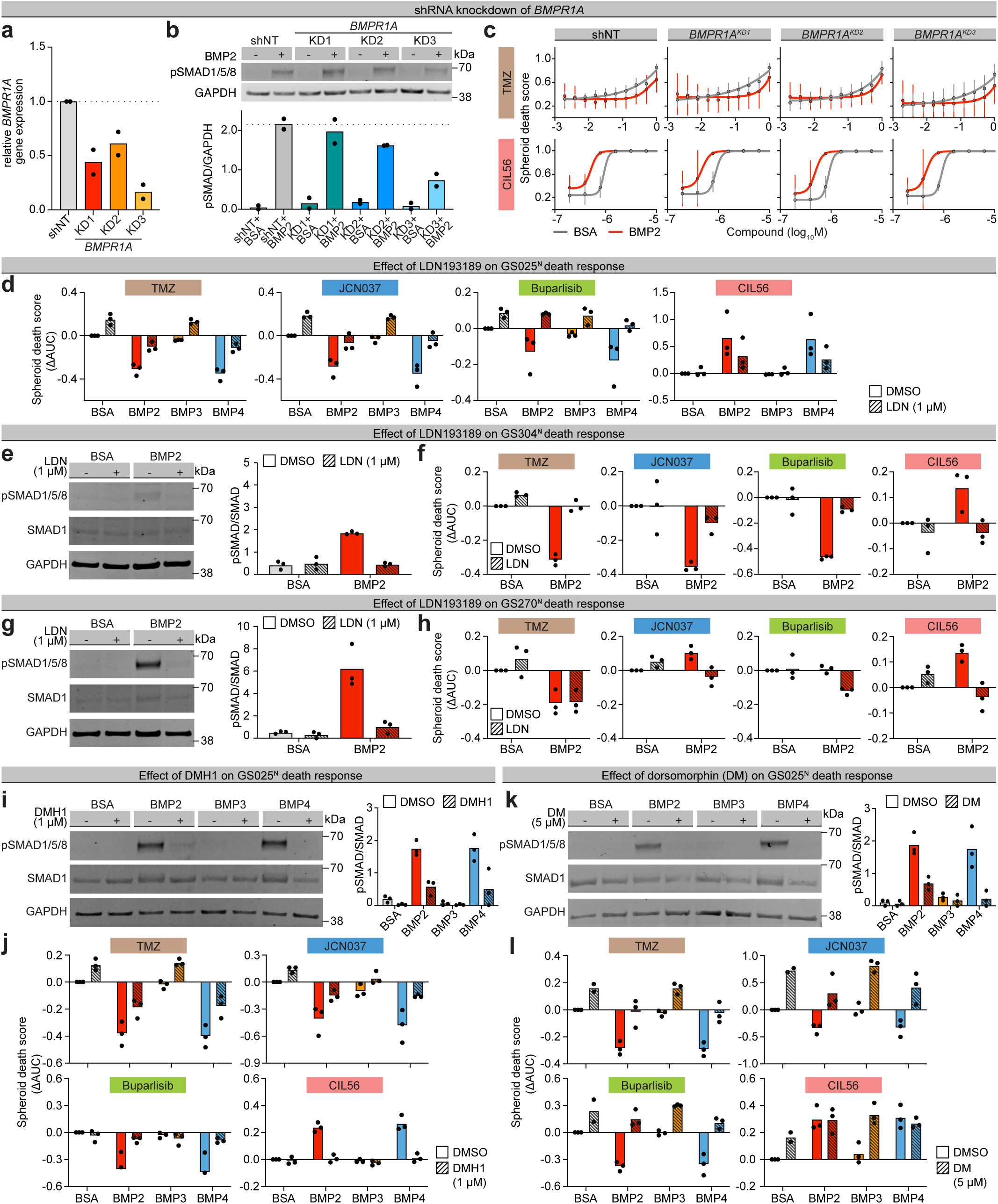
BMPR1 inhibition reverses BMP-mediated cell death effects. a, RT-qPCR validation of *BMPR1A* knockdown. b, Immunoblot analysis of SMAD1/5/8 activation (pSMAD1/5/8) following *BMPR1A* knockdown. c, Effect of *BMPR1A* knockdown on temozolomide (TMZ)- and CIL56-induced death. d, Effect of BMPR1A and BMPR1B pharmacological inhibition with LDN193189 (LDN; 1 µM) on BMP2-mediated modulation of death in GS025^N^ gliomaspheres. e, Immunoblot analysis of SMAD1/5/8 activation following LDN treatment in GS304^N^ gliomaspheres. f, Effect of LDN on BMP2-mediated modulation of death in GS304^N^ gliomaspheres. g, Immunoblot analysis of SMAD1/5/8 activation following LDN treatment in GS270^N^ gliomaspheres. h, Effect of LDN on BMP2-mediated modulation of death in GS270^N^ gliomaspheres. i, Immunoblot analysis of SMAD1/5/8 activation following pharmacological inhibition of BMPR1A and BMPR1B with DMH1 (1 µM) in GS025^N^ gliomaspheres. j, Effect of DMH1 on BMP2-mediated modulation of death in GS025^N^ gliomaspheres. k, Immunoblot analysis of SMAD1/5/8 activation following pharmacological inhibition of BMPR1A and BMPR1B with dorsomorphin (DM; 5 µM) in GS025^N^ gliomaspheres. l, Effect of DM on BMP2-mediated modulation of death in GS025^N^ gliomaspheres. *BMPR1A* shRNA knockdown experiments (a), (b), and (c) consist of two independent experimental replicates; all others consist of three independent experiments.

**Extended Data Figure 5.**
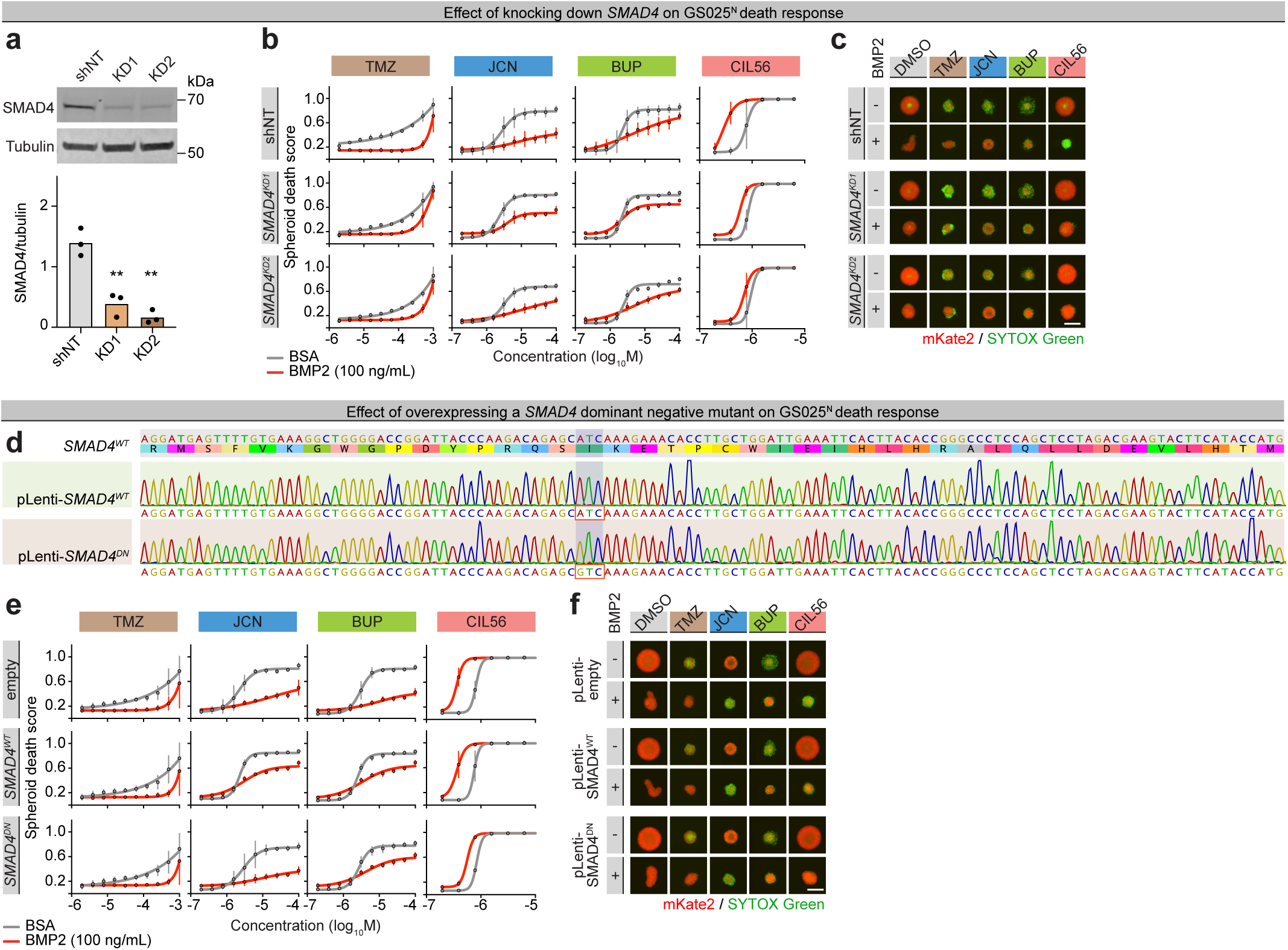
Partial loss of SMAD4 suppresses BMP2 modulation of CIL56 sensitivity. a, Immunoblot validation of *SMAD4* knockdown. b, Effect of shRNA-mediated knockdown of *SMAD4* on BMP2-modulation of spheroid death. c, Representative images corresponding to panel b. d, Sequencing traces confirming the nucleotide change encoding the *SMAD4* I500V dominant negative mutation. e, Effect of overexpressing wildtype and dominant negative *SMAD4* on BMP2-mediated modulation of spheroid death. f, Representative images corresponding to panel e. Data in (b) and (e) represent mean ± s.d. from three independent experiments. Immunoblots and gliomasphere images are representative of three independent experiments. Scale bars = 400 µm.

**Extended Data Figure 6.**
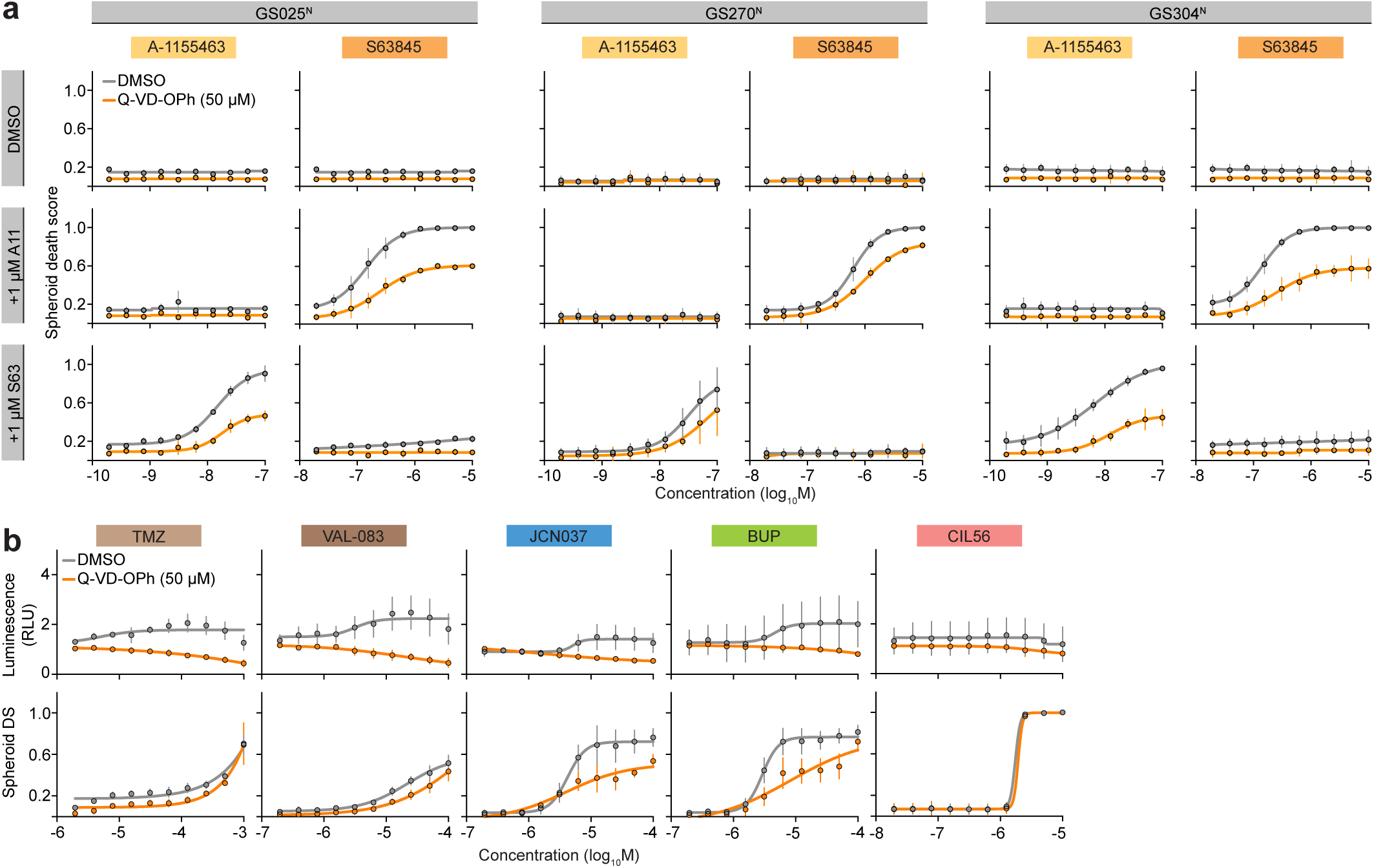
Caspase-dependence of compounds tested. a, Dose response curves of A-1155463 and S63845 in three patient-derived gliomaspheres, with and without the caspase inhibitor Q-VD-OPh (50 µM). b, Caspase-Glo 3/7 3D and HADES analysis showing compound induced death despite suppression of caspase activity. Data represent mean ± s.d. from three independent experiments. All ligand treatments = 100 ng/mL.

**Extended Data Figure 7:**
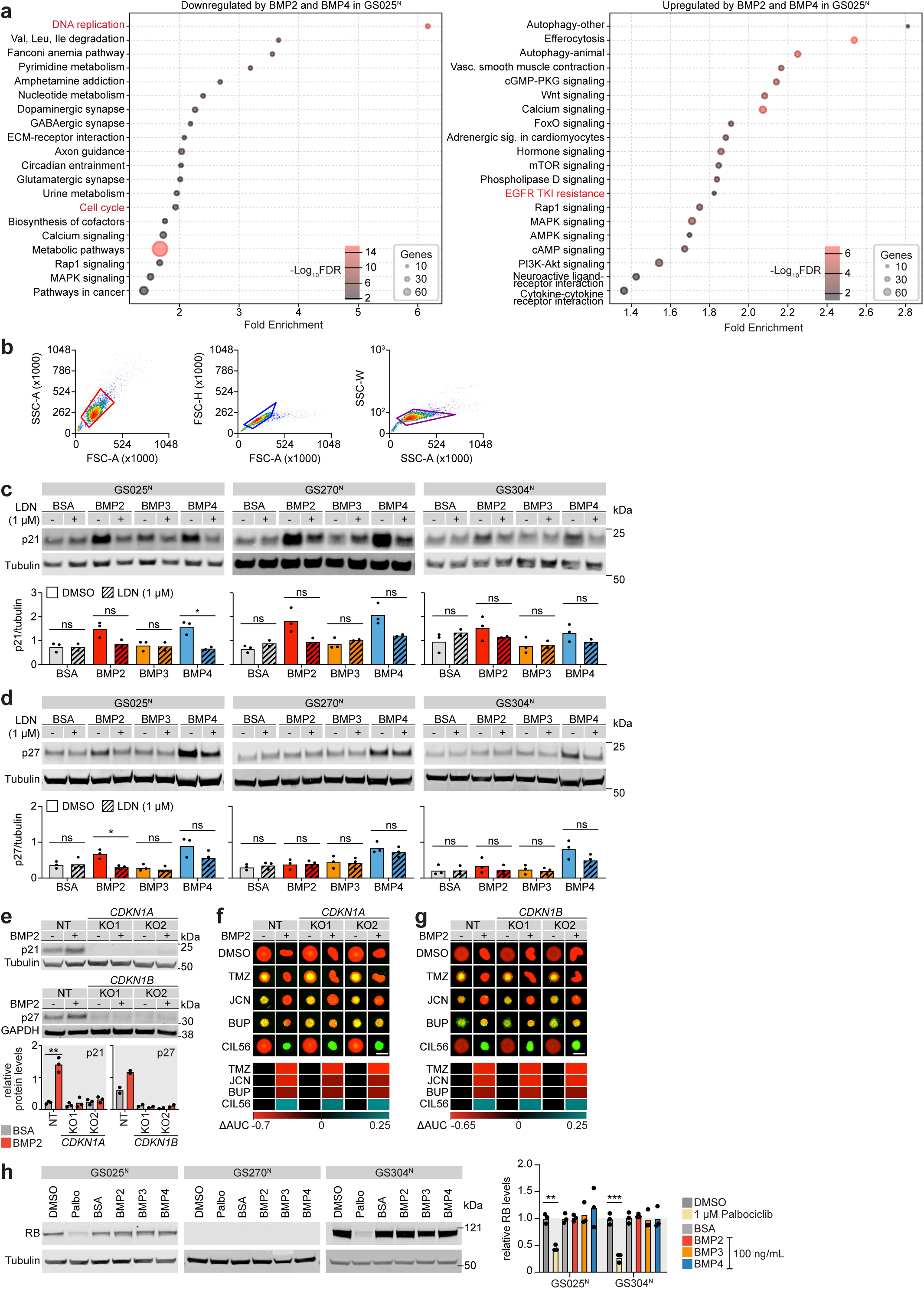
BMP effects on spheroid growth and CDK inhibitors. a, KEGG pathway analysis of GS025^N^ spheroids treated with BMP2 or BMP4 b, Gating strategy used in Figure 4j,k. SSC x FSC signals were used to gate for typical GBM populations, FSC-H x FSC-A and SSC-W x SSC-A signals were used to exclude doublets. Representative plots show data collected with GS025^N^ cells. c, Immunoblots showing the effects of BMP2/4 on p21 abundance across three patient-derived gliomasphere lines. d, Immunoblots showing the effects of BMP2/4 on p27 abundance across three patient-derived gliomasphere lines. e, Immunoblot validation of *CDKN1A* and *CDKN1B* rCRISPR targeting. f, Effect of rCRISPR-mediated disruption of *CDKN1A* on GS025^R^ gliomasphere death. g, Effect of rCRISPR-mediated disruption of *CDKN1B* on GS025^R^ gliomasphere death. h, Immunoblots showing the effects of palbociclib (Palbo; 1 µM) on RB levels. All data besides *CDKN1B* targeting experiments (in e and g) consist of three independent experiments. *CDKN1B* targeting experiments consist of two independent experiments. For panels (f) and (g), TMZ: temozolomide was used at 500 µM, JCN: JCN037 was used at 12.5 µM, BUP: buparlisib was used at 12.5 µM, and CIL56 was used at 1.25 µM. Gliomasphere death was summarized as heatmaps in the lower sub-panels. ΔAUC values= AUC^ligand^ - AUC^BSA^. Red indicates decreased spheroid death, blue indicates increased spheroid death. All ligand treatments = 100 ng/mL. Scale bar = 400 µm.

**Extended Data Figure 8.**
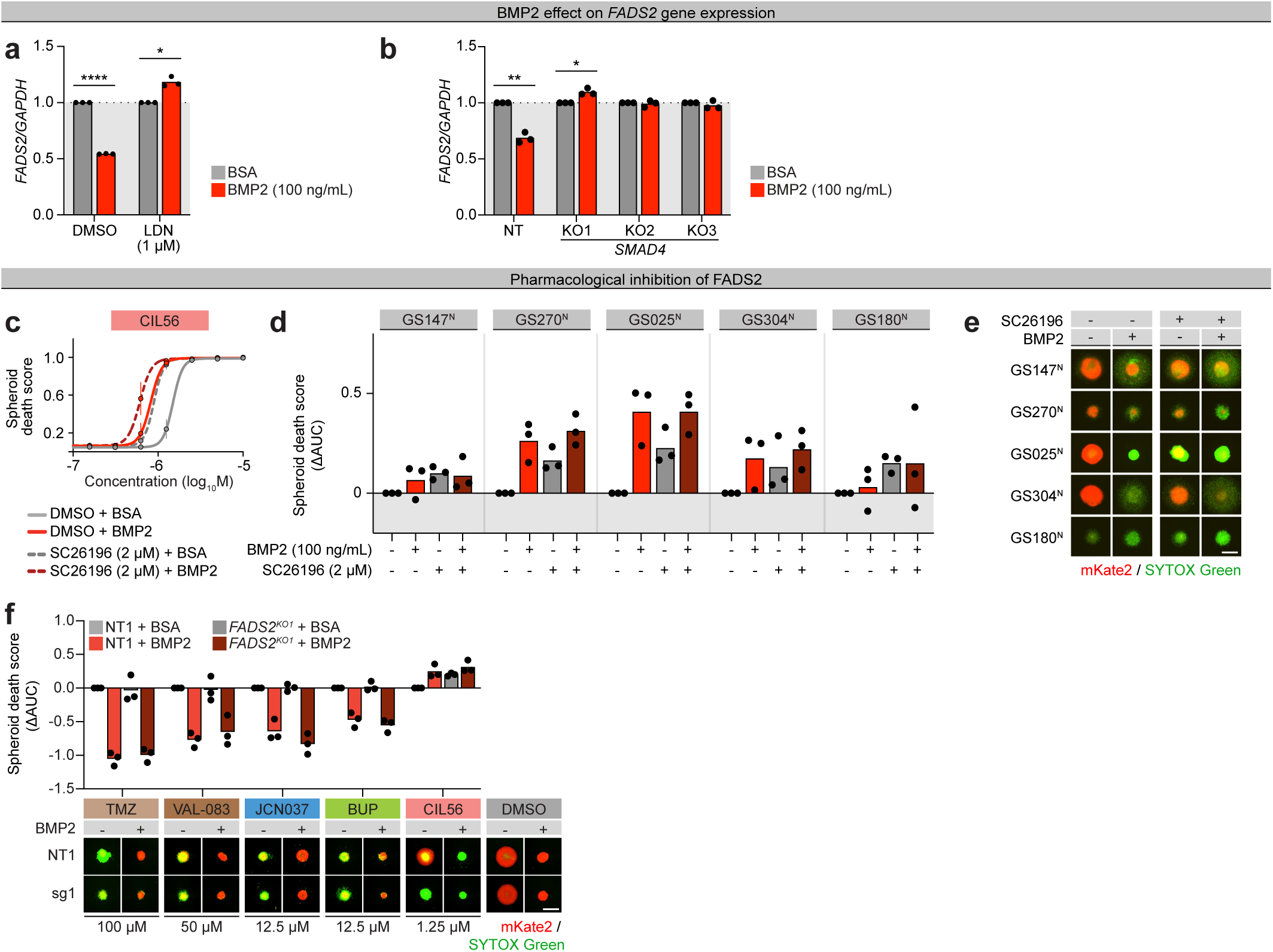
BMP2 increase LiDN sensitivity by decreasing *FADS2* expression. a, Effect of BMPR1A and BMPR1B pharmacological inhibition with LDN193189 (LDN; 1 µM) on BMP2-mediated downregulation of *FADS2* in GS025^N^ gliomaspheres as determined by RT-qPCR. b, Effect of *SMAD4* gene disruption on BMP2-mediated downregulation of *FADS2* in GS025^N^ gliomaspheres as determined by RT-qPCR. c, Effect of FADS2 pharmacological inhibition with SC26196 (2 µM) on CIL56-induced death in GS025^N^ gliomaspheres. d, Effect of FADS2 pharmacological inhibition with SC26196 (2 µM) on CIL56-induced death in five patient-derived gliomasphere lines. ΔAUC = AUC_treated_ - AUC_BSA+DMSO_. e, Representative images corresponding to panel d. f, Effect of rCRISPR-mediated gene disruption of *FADS2* on LiDN and non-LiDN death in GS025^R^ gliomaspheres. ΔAUC = AUC_treated_ - AUC_NT+BSA+DMSO_. Data in (a), (b), (d), and (f) represent mean of three independent experimental replicates. Data in (c) represent mean ± s.d. from three independent experiments. Images in (e) and (f) are representative of three independent experiments. All ligand treatments = 100 ng/mL. Scale bars = 400 µm.

**Extended Data Figure 9.**
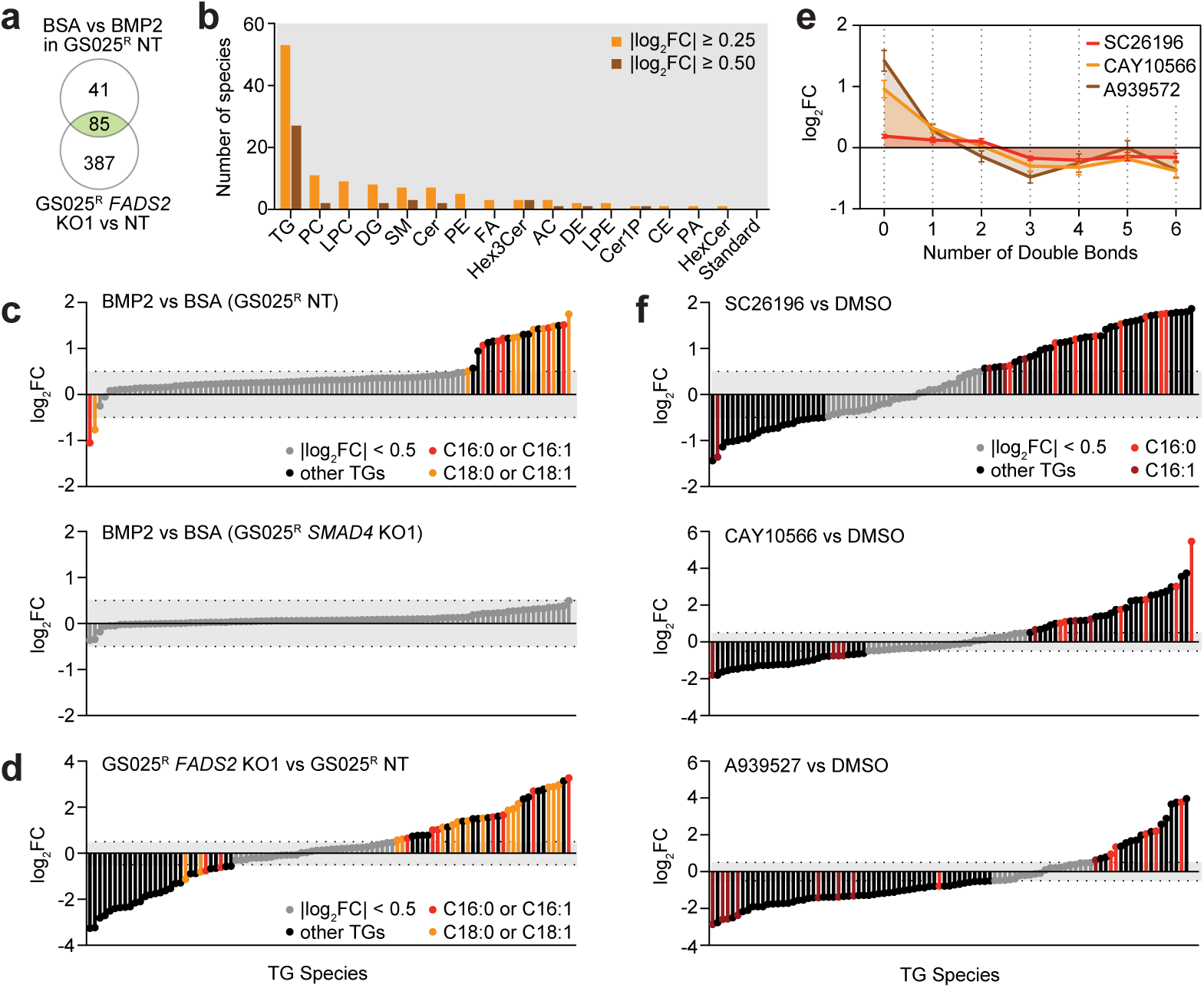
Lipid rewiring associated with BMP2 signaling and *FADS2* disruption. a, Venn diagram highlighting lipid species commonly increased (|log2FC| ≥ 0.5) by BMP2 treatment (100 ng/mL) and *FADS2* rCRISPR targeting. b, Number of lipid species changed in the same direction by BMP2 and *FADS2* rCRISPR targeting, grouped by lipid class. c, Triglyceride species altered by BMP2 treatment (100 ng/mL) in GS025^R^ non-targeting (NT) and *SMAD4* disrupted cells. d, Triglyceride species altered following *FADS2* disruption. e, Effects of SC26196, CAY10566, and A939572 (2 µM) on global lipid saturation. f, Triglyceride species altered by SC26196, CAY10566, and A939572 (2 µM) in GS025^R^ NT cells.

## REFERENCES

1. Omuro, A. & DeAngelis, L.M. Glioblastoma and other malignant gliomas: a clinical review. JAMA 310, 1842–1850 (2013).

2. Stupp, R. et al. Radiotherapy plus concomitant and adjuvant temozolomide for glioblastoma. N Engl J Med 352, 987–996 (2005).

3. Singh, N., Miner, A., Hennis, L. & Mittal, S. Mechanisms of temozolomide resistance in glioblastoma - a comprehensive review. Cancer Drug Resist 4, 17–43 (2021).

4. Ochs, K. & Kaina, B. Apoptosis induced by DNA damage O6-methylguanine is Bcl-2 and caspase-9/3 regulated and Fas/caspase-8 independent. Cancer Res 60, 5815–5824 (2000).

5. Fernandez, E.G. et al. Integrated molecular and functional characterization of the intrinsic apoptotic machinery identifies therapeutic vulnerabilities in glioma. Nat Commun 15, 10089 (2024).

6. Dewdney, B. et al. From signalling pathways to targeted therapies: unravelling glioblastoma’s secrets and harnessing two decades of progress. Signal Transduct Target Ther 8, 400 (2023).

7. Wen, P.Y. et al. Buparlisib in Patients With Recurrent Glioblastoma Harboring Phosphatidylinositol 3-Kinase Pathway Activation: An Open-Label, Multicenter, Multi-Arm, Phase II Trial. J Clin Oncol 37, 741–750 (2019).

8. Koul, D. et al. Antitumor activity of NVP-BKM120--a selective pan class I PI3 kinase inhibitor showed differential forms of cell death based on p53 status of glioma cells. Clin Cancer Res 18, 184–195 (2012).

9. Chagoya, G. et al. Efficacy of osimertinib against EGFRvIII+ glioblastoma. Oncotarget 11, 2074–2082 (2020).

10. Tsang, J.E. et al. Development of a Potent Brain-Penetrant EGFR Tyrosine Kinase Inhibitor against Malignant Brain Tumors. ACS Med Chem Lett 11, 1799–1809 (2020).

11. Wen, P.Y. et al. First-in-Human Phase I Study to Evaluate the Brain-Penetrant PI3K/mTOR Inhibitor GDC-0084 in Patients with Progressive or Recurrent High-Grade Glioma. Clin Cancer Res 26, 1820–1828 (2020).

12. Cheng, Y.L. et al. Multiplexed single-cell lineage tracing of mitotic kinesin inhibitor resistance in glioblastoma. Cell Rep 43, 114139 (2024).

13. Strik, H. et al. BCL-2 family protein expression in initial and recurrent glioblastomas: modulation by radiochemotherapy. J Neurol Neurosurg Psychiatry 67, 763–768 (1999).

14. Koessinger, A.L. et al. Increased apoptotic sensitivity of glioblastoma enables therapeutic targeting by BH3-mimetics. Cell Death Differ 29, 2089–2104 (2022).

15. Minami, J.K. et al. CDKN2A deletion remodels lipid metabolism to prime glioblastoma for ferroptosis. Cancer Cell 41, 1048–1060 e1049 (2023).

16. Lee, J. et al. Tumor stem cells derived from glioblastomas cultured in bFGF and EGF more closely mirror the phenotype and genotype of primary tumors than do serum-cultured cell lines. Cancer Cell 9, 391–403 (2006).

17. Qin, E.Y. et al. Neural Precursor-Derived Pleiotrophin Mediates Subventricular Zone Invasion by Glioma. Cell 170, 845–859 e819 (2017).

18. Dixon, S.J. & Lee, M.J. Quick tips for interpreting cell death experiments. Nat Cell Biol 25, 1720–1723 (2023).

19. Forcina, G.C., Conlon, M., Wells, A., Cao, J.Y. & Dixon, S.J. Systematic Quantification of Population Cell Death Kinetics in Mammalian Cells. Cell Syst 4, 600–610 e606 (2017).

20. Galluzzi, L. et al. Molecular mechanisms of cell death: recommendations of the Nomenclature Committee on Cell Death 2018. Cell Death Differ 25, 486–541 (2018).

21. Venkatesh, H.S. et al. Neuronal Activity Promotes Glioma Growth through Neuroligin-3 Secretion. Cell 161, 803–816 (2015).

22. Hara, T. et al. Interactions between cancer cells and immune cells drive transitions to mesenchymal-like states in glioblastoma. Cancer Cell 39, 779–792 e711 (2021).

23. Wilson, T.R. et al. Widespread potential for growth-factor-driven resistance to anticancer kinase inhibitors. Nature 487, 505–509 (2012).

24. Ko, P.J. et al. A ZDHHC5-GOLGA7 Protein Acyltransferase Complex Promotes Nonapoptotic Cell Death. Cell Chem Biol 26, 1716–1724 e1719 (2019).

25. Leak, L. et al. Tegavivint triggers TECR-dependent nonapoptotic cancer cell death. Nat Chem Biol 21, 1873–1884 (2025).

26. Fontebasso, A.M. et al. Recurrent somatic mutations in ACVR1 in pediatric midline high-grade astrocytoma. Nat Genet 46, 462–466 (2014).

27. Taylor, K.R. et al. Recurrent activating ACVR1 mutations in diffuse intrinsic pontine glioma. Nat Genet 46, 457–461 (2014).

28. Wu, G. et al. The genomic landscape of diffuse intrinsic pontine glioma and pediatric non-brainstem high-grade glioma. Nat Genet 46, 444–450 (2014).

29. Buczkowicz, P. et al. Genomic analysis of diffuse intrinsic pontine gliomas identifies three molecular subgroups and recurrent activating ACVR1 mutations. Nat Genet 46, 451–456 (2014).

30. Jacob, F. et al. A Patient-Derived Glioblastoma Organoid Model and Biobank Recapitulates Inter- and Intra-tumoral Heterogeneity. Cell 180, 188–204 e122 (2020).

31. Neftel, C. et al. An Integrative Model of Cellular States, Plasticity, and Genetics for Glioblastoma. Cell 178, 835–849 e821 (2019).

32. Tao, Z.F. et al. Discovery of a Potent and Selective BCL-XL Inhibitor with in Vivo Activity. ACS Med Chem Lett 5, 1088–1093 (2014).

33. Caserta, T.M., Smith, A.N., Gultice, A.D., Reedy, M.A. & Brown, T.L. Q-VD-OPh, a broad spectrum caspase inhibitor with potent antiapoptotic properties. Apoptosis 8, 345–352 (2003).

34. Mishra, J., Dickinson, J.D. & Maurya, S.K. Brain microenvironment orchestrates highly aggressive tumor variants: current trends and therapeutic approaches. Front Aging Neurosci 17, 1666837 (2025).

35. Kotschy, A. et al. The MCL1 inhibitor S63845 is tolerable and effective in diverse cancer models. Nature 538, 477–482 (2016).

36. Dixon, S.J. et al. Ferroptosis: an iron-dependent form of nonapoptotic cell death. Cell 149, 1060–1072 (2012).

37. Stupp, R. et al. Radiotherapy plus concomitant and adjuvant temozolomide for glioblastoma. N Engl J Med 352, 987–996 (2005).

38. Guo, C., et al. VAL-083 is effective in patients with newly-diagnosed MGMT-unmethylated glioblastoma: report of phase II study. Discov Oncol (2025).

39. Wen, P.Y. et al. Buparlisib in Patients With Recurrent Glioblastoma Harboring Phosphatidylinositol 3-Kinase Pathway Activation: An Open-Label, Multicenter, Multi-Arm, Phase II Trial. J Clin Oncol 37, 741–750 (2019).

40. Fuentes-Baile, M. et al. Differential Effects of IGF-1R Small Molecule Tyrosine Kinase Inhibitors BMS-754807 and OSI-906 on Human Cancer Cell Lines. Cancers (Basel) 12 (2020).

41. Lombardi, G. et al. Regorafenib compared with lomustine in patients with relapsed glioblastoma (REGOMA): a multicentre, open-label, randomised, controlled, phase 2 trial. Lancet Oncol 20, 110–119 (2019).

42. Roth, P. et al. Marizomib for patients with newly diagnosed glioblastoma: A randomized phase 3 trial. Neuro Oncol 26, 1670–1682 (2024).

43. Jin, S. et al. Inference and analysis of cell-cell communication using CellChat. Nat Commun 12, 1088 (2021).

44. Noel, F. et al. Dissection of intercellular communication using the transcriptome-based framework ICELLNET. Nat Commun 12, 1089 (2021).

45. Hou, R., Denisenko, E., Ong, H.T., Ramilowski, J.A. & Forrest, A.R.R. Predicting cell-to-cell communication networks using NATMI. Nat Commun 11, 5011 (2020).

46. Gao, W. et al. Targeting mesenchymal monocyte-derived macrophages to enhance the sensitivity of glioblastoma to temozolomide by inhibiting TNF/CELSR2/p65/Kla-HDAC1/EPAS1 axis. J Adv Res 80, 925–941 (2026).

47. Murakami, M., Kamimura, D. & Hirano, T. Pleiotropy and Specificity: Insights from the Interleukin 6 Family of Cytokines. Immunity 50, 812–831 (2019).

48. Chen, D., Zhao, M. & Mundy, G.R. Bone morphogenetic proteins. Growth Factors 22, 233–241 (2004).

49. Al-Dalahmah, O. et al. Re-convolving the compositional landscape of primary and recurrent glioblastoma reveals prognostic and targetable tissue states. Nat Commun 14, 2586 (2023).

50. Cuny, G.D. et al. Structure-activity relationship study of bone morphogenetic protein (BMP) signaling inhibitors. Bioorg Med Chem Lett 18, 4388–4392 (2008).

51. Hao, J. et al. In vivo structure-activity relationship study of dorsomorphin analogues identifies selective VEGF and BMP inhibitors. ACS Chem Biol 5, 245–253 (2010).

52. Yu, P.B. et al. Dorsomorphin inhibits BMP signals required for embryogenesis and iron metabolism. Nat Chem Biol 4, 33–41 (2008).

53. Ehata, S. & Miyazono, K. Bone Morphogenetic Protein Signaling in Cancer; Some Topics in the Recent 10 Years. Front Cell Dev Biol 10, 883523 (2022).

54. Alankarage, D. et al. Myhre syndrome is caused by dominant-negative dysregulation of SMAD4 and other co-factors. Differentiation 128, 1–12 (2022).

55. Hendel, A. et al. Chemically modified guide RNAs enhance CRISPR-Cas genome editing in human primary cells. Nat Biotechnol 33, 985–989 (2015).

56. Potter, D.S., Du, R., Bhola, P., Bueno, R. & Letai, A. Dynamic BH3 profiling identifies active BH3 mimetic combinations in non-small cell lung cancer. Cell Death Dis 12, 741 (2021).

57. Piccirillo, S.G. et al. Bone morphogenetic proteins inhibit the tumorigenic potential of human brain tumour-initiating cells. Nature 444, 761–765 (2006).

58. Chang, S.F. et al. BMP-4 induction of arrest and differentiation of osteoblast-like cells via p21 CIP1 and p27 KIP1 regulation. Mol Endocrinol 23, 1827–1838 (2009).

59. Sachdeva, R. et al. BMP signaling mediates glioma stem cell quiescence and confers treatment resistance in glioblastoma. Sci Rep 9, 14569 (2019).

60. Finn, R.S. et al. PD 0332991, a selective cyclin D kinase 4/6 inhibitor, preferentially inhibits proliferation of luminal estrogen receptor-positive human breast cancer cell lines in vitro. Breast Cancer Res 11, R77 (2009).

61. Goel, S., Bergholz, J.S. & Zhao, J.J. Targeting CDK4 and CDK6 in cancer. Nat Rev Cancer 22, 356–372 (2022).

62. Triki, M. et al. mTOR Signaling and SREBP Activity Increase FADS2 Expression and Can Activate Sapienate Biosynthesis. Cell Rep 31, 107806 (2020).

63. Hasegawa, K. et al. Delta-6 desaturase FADS2 is a tumor-promoting factor in cholangiocarcinoma. Cancer Sci 115, 3346–3357 (2024).

64. Obukowicz, M.G. et al. Novel, selective delta6 or delta5 fatty acid desaturase inhibitors as antiinflammatory agents in mice. J Pharmacol Exp Ther 287, 157–166 (1998).

65. Wunderling, K., Zurkovic, J., Zink, F., Kuerschner, L. & Thiele, C. Triglyceride cycling enables modification of stored fatty acids. Nat Metab 5, 699–709 (2023).

66. Zhou, K. et al. A new glioma grading model based on histopathology and Bone Morphogenetic Protein 2 mRNA expression. Sci Rep 10, 18420 (2020).

67. Bao, Z. et al. BMP4, a strong better prognosis predictor, has a subtype preference and cell development association in gliomas. J Transl Med 11, 100 (2013).

68. Bos, E.M. et al. Local delivery of hrBMP4 as an anticancer therapy in patients with recurrent glioblastoma: a first-in-human phase 1 dose escalation trial. Mol Cancer 22, 129 (2023).

69. Murray, M.B., Leak, L.B., Lee, W.C. & Dixon, S.J. Protocol for detection of ferroptosis in cultured cells. STAR Protoc 4, 102457 (2023).

70. Kim, K.H. & Sederstrom, J.M. Assaying Cell Cycle Status Using Flow Cytometry. Curr Protoc Mol Biol 111, 28 26 21-28 26 11 (2015).

71. Hughes, C.S. et al. Single-pot, solid-phase-enhanced sample preparation for proteomics experiments. Nat Protoc 14, 68–85 (2019).

72. Demichev, V., Messner, C.B., Vernardis, S.I., Lilley, K.S. & Ralser, M. DIA-NN: neural networks and interference correction enable deep proteome coverage in high throughput. Nat Methods 17, 41–44 (2020).

